# Both major xanthophyll cycles present in nature can provide Non-Photochemical Quenching in the model diatom *Phaeodactylum tricornutum*

**DOI:** 10.1101/2024.03.19.584964

**Authors:** Chiara E. Giossi, Marie A. Wünsch, Oliver Dautermann, Alexander F. Schober, Jochen M. Buck, Peter G. Kroth, Martin Lohr, Bernard Lepetit

## Abstract

Xanthophyll cycling contributes to photoprotection by regulating Non-Photochemical Quenching (NPQ). While most photosynthetic eukaryotes including land plants use the violaxanthin cycle, some algae like diatoms and haptophytes rely on the diadinoxanthin cycle for photoprotection. These algae also contain minor amounts of violaxanthin cycle pigments, serving as precursors in xanthophylls biosynthesis. Both cycles are catalyzed by the enzymes violaxanthin de-epoxidase (VDE) and zeaxanthin epoxidase (ZEP). Here, we characterized the role of *VDE* and different ZEP-encoding paralogs (*ZEP2* and *ZEP3*) in the model diatom *Phaeodactylum tricornutum*. While knockout of *VDE* and *ZEP3* significantly impaired the diadinoxanthin cycle, lack of *ZEP2* led to sustained accumulation of the violaxanthin cycle instead of diadinoxanthin cycle pigments under high irradiance, with no negative effect on NPQ capacity. We demonstrate that both major xanthophyll cycles present in nature can function with comparable efficiency within the same species, offering a new perspective on the evolution of xanthophyll-mediated photoprotection.

## Introduction

Diatoms are amongst the main primary producers on Earth (Field et al., 1998; Falkowski et al., 2004), dominating phytoplankton and microphytobenthic communities in nutrient-rich turbulent waters where irradiance can change very rapidly due to daily fluctuations and to the movement of water and sediments (Tozzi et al., 2004; Lavaud, 2007). To cope with this highly dynamic light stress and avoid the formation of reactive oxygen species and consequent photoinhibition, diatoms have evolved different photoprotection strategies (Lavaud et al., 2007; Kan et al., 2023). One of these is Non-Photochemical Quenching (NPQ), a ubiquitous multi-component mechanism that dissipates the surplus of excitation energy as heat, thus preventing the over-saturation of the photosynthetic apparatus (Takahashi and Badger, 2011; Domingues et al., 2012; Goss and Lepetit, 2015; Bassi and Dall’Osto, 2021). qE, or high energy-state quenching, is fastest component of NPQ, taking place in the light-harvesting complexes of photosystem II (PSII) (Takahashi and Badger, 2011; Domingues et al., 2012; Goss and Lepetit, 2015; Bassi and Dall’Osto, 2021). In diatoms, this mechanism relies on antenna proteins of the Lhcx family (Bailleul et al., 2010; Lepetit et al., 2017; Buck et al., 2019; Buck et al., 2021).

In most photosynthetic eukaryotes, the on- and offset of qE is regulated by the xanthophyll cycle, a rapid interconversion of carotenoids that relies on removal or addition of epoxy groups (Latowski et al., 2011). Two major xanthophyll cycles exist in nature (Latowski et al., 2011; Goss and Lepetit, 2015): i) the violaxanthin (Vx) cycle, a two-step reaction involving Vx, antheraxanthin (Ax) and zeaxanthin (Zx), present in green algae, land plants and several classes of heterokont algae; and ii) the diadinoxanthin (Dd) cycle, responsible for the interconversion of only two pigments, Dd and diatoxanthin (Dt), found in haptophytes and other heterokonts (including diatoms). Both cycles are tightly regulated via light-dependent changes in the trans-thylakoidal ΔpH (Latowski et al., 2011; Goss and Lepetit, 2015). In diatoms, the light induced ΔpH activates the enzyme violaxanthin de-epoxidase (VDE), converting Dd into Dt and thus triggering qE. When light stress and ΔpH are relaxed, Dt is converted back to Dd by the enzyme zeaxanthin epoxidase (ZEP), completing the cycle and thus abolishing qE (Goss et al., 2006a; Blommaert et al., 2021). Similarly, in plants and algae displaying the Vx cycle qE is triggered by the de-epoxidation of Vx to Zx through the intermediate Ax, and Vx is restored by the re-epoxidation of Zx (Latowski et al., 2011). Interestingly, diatoms and other algae using the Dd cycle can accumulate pigments of the Vx cycle under extreme irradiance (Lohr and Wilhelm, 1999; Lohr and Wilhelm, 2001; Kuczynska et al., 2020). This was explained by the essential role of Vx as intermediate in the biosynthesis of Dd and fucoxanthin (Lohr, 2011; Dambek et al., 2012). However, it is unclear whether Zx and Ax can also trigger NPQ in diatoms. Diatoms possess multiple paralogs encoding for potential xanthophyll cycle enzymes, originated via gene duplication (Coesel et al., 2008; Frommolt et al., 2008). In the model diatom *Phaeodactylum tricornutum*, the role of ZEP1, VDL1 and VDL2 in the fucoxanthin biosynthetic pathway has been recently elucidated (Dautermann et al., 2020; Bai et al., 2022). Hence, the photoprotective function must be provided by other paralogs such as *VDE*, *VDR, ZEP2* and/or *ZEP3*. Indeed, RNA silencing (Lavaud et al., 2012) and recombinant protein experiments (Bojko et al., 2013; Olchawa-Pajor et al., 2019) pointed to VDE as the major regulator of Dd de-epoxidation in *P. tricornutum*. Moreover, *VDE* and *ZEP3* are arranged in a gene cluster conserved in *P. tricornutum* and other diatom lineages, suggesting shared evolutionary history and conserved functions (Frommolt et al., 2008; Bai et al., 2022). Thus, VDE and ZEP3 could be the main regulators of the Dd cycle in *P. tricornutum*. On the contrary, ZEP2 may have an essential role in other photoprotection mechanisms and/or carotenoid biosynthesis (Wilhelm et al., 2006; Frommolt et al., 2008; Kuczynska et al., 2020).

In this work, we investigated the role of three xanthophyll cycle enzymes (ZEP2, ZEP3 and VDE) in the Dd and Vx cycle in *P. tricornutum* through the respective knockout mutants. Our results highlight VDE and ZEP3 as major drivers of the Dd cycle: the VDE-deficient mutant (*vde KO*) was completely Dd cycle-deficient, while the ZEP3-deficient mutant (*zep3 KO*) did not rapidly re-epoxidize Dt after light stress. ZEP2-deficient mutants (*zep2 KO*) accumulated significantly large amounts of Zx and Ax under high light, at the expense of Dt; nevertheless, their NPQ capacity was comparable to the *wild type* (wt). This proves for the first time that both major xanthophyll cycles present in nature can provide NPQ in the same organism and suggests that Dt and Zx/Ax can contribute to the on- and offset of qE with comparable efficiency in diatoms.

## Results

### Knockouts of VDE and ZEP3 impair photoprotective xanthophyll cycling, while lack of ZEP2 enhances the accumulation of Vx cycle pigments in response to high light (HL)

We generated knockout mutants for *ZEP2*, *ZEP3* and *VDE* throug CRISPR-Cas 9 approach (**Supplemental Note S1**). When grown under continuous low light (LL), no major differences in pigment content were observed between all newly generated mutant lines and wt (**Supplemental Table S2**).

However, after 5 days of 6 h HL:18 h LL regime each mutant displayed differences in xanthophyll cycling when compared to wt (**Fig. 1, Supplemental Fig. S5-S7**). In the wt, Dd slightly decreased after 6h of HL, concomitant with a massive Dt increase. In addition, minor amounts of Zx accumulated. During the following 30 min of recovery in LL, Dt and Zx were almost entirely epoxidized back to Dd and Vx. As expected, *vde KO* did not accumulate Dt or Zx during 6h of HL, but instead showed a strong increase of Dd and a small but statistically significant increase in Vx. Conversely, *zep3 KO* accumulated the highest amount of Dt under HL but was not able to convert it back to Dd during LL recovery. Finally, *zep2 KO* displayed the Dd cycle in response to changing light, but these pigments were present in significantly lower amounts compared to wt. Instead, this mutant was accumulating and cycling Vx cycle pigments to a much higher extent than all other strains: Zx accumulated in response to 6h of HL and then decreased during the early stages of recovery (5 min of LL), while a transient pool of Ax accumulated; after 30min of LL, Vx accumulated at the expense of both Zx and Ax.

**Fig. 1.**
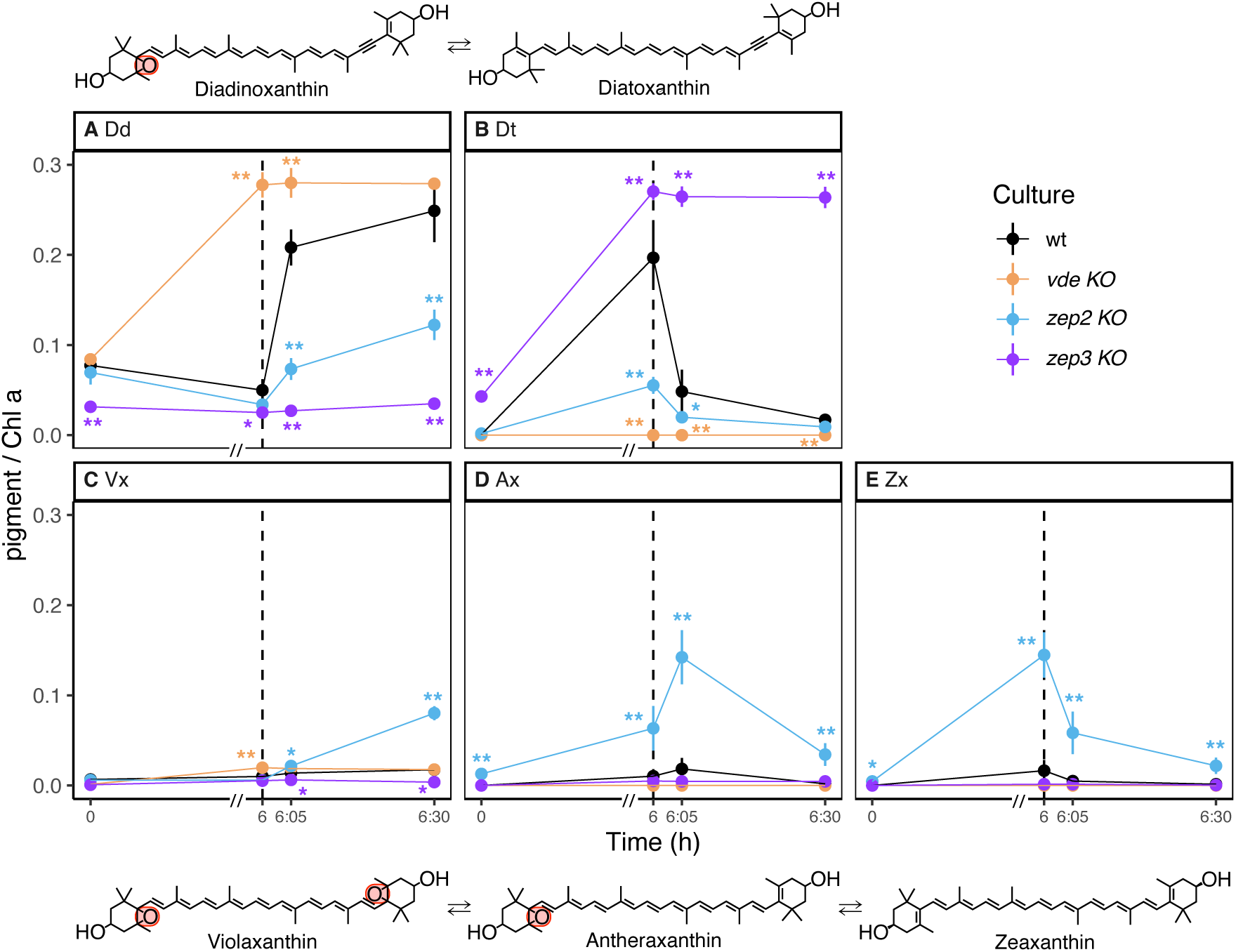
Content of Dd and Vx cycle pigments in wt, *vde KO*, *zep2 KO* and *zep3 KO* after 5 days of HL:LL regime (average ± sd, n=3). Pigment content is expressed as pigment:chlorophyll *a* ratio (mol/mol). Time 0 is defined as the beginning of the HL phase (i.e. the sampling point in LL that immediately precedes the onset of HL on the fifth day of HL:LL treatment). The 6 h of HL phase and 30 min of recovery phase (LL) are separated with a dashed line. On the x-axis, recovery phase (from 6 to 6:30 h) was artificially enlarged to allow better visualization; axis break is indicated by a double dash (//). (*A*) diadinoxanthin; (*B*) diatoxanthin; (*C*) violaxanthin; (*D*) antheraxanthin; (*E*) zeaxanthin. Epoxy groups are highlighted with a red rectangle. Statistical significance marks indicate significant differences between the corresponding mutant line and wt at each time point, according to adjusted p-value of multiple comparison t-test (*: p < 0.05; **: p < 0.005).

The phenotypes described for all mutants were reverted to wt in mutant lines complemented with the respective native genes under native promoters (**Supplemental Fig. S8-10**).

### *De novo* synthesis drives the light-induced accumulation of photoprotective xanthophylls in all mutants, with no effect on the total pigment pool

After 6 h of HL, *vde KO* and *zep3 KO* displayed no major difference in total Dd+Dt pool compared to wt (**Fig. 2A**), and in all three lines the Vx cycle pool was almost absent (**Fig. 2B**). In contrast, *zep2 KO* displayed a much lower Dd+Dt content (about 20% of the total xanthophyll cycle pigments, **Fig. 2A**) and instead accumulated a massive total pool of Vx+Ax+Zx (about 80% of the total, **Fig. 2B**). All investigated lines approximatively tripled their total xanthophyll cycle pigment content, with no significant difference between each mutant and wt (**Fig. 2C**). This suggests that the rate of de-novo biosynthesis of carotenoids is not affected by any of the target mutations, thus being likely regulated in the same way in all investigated lines.

**Fig. 2.**
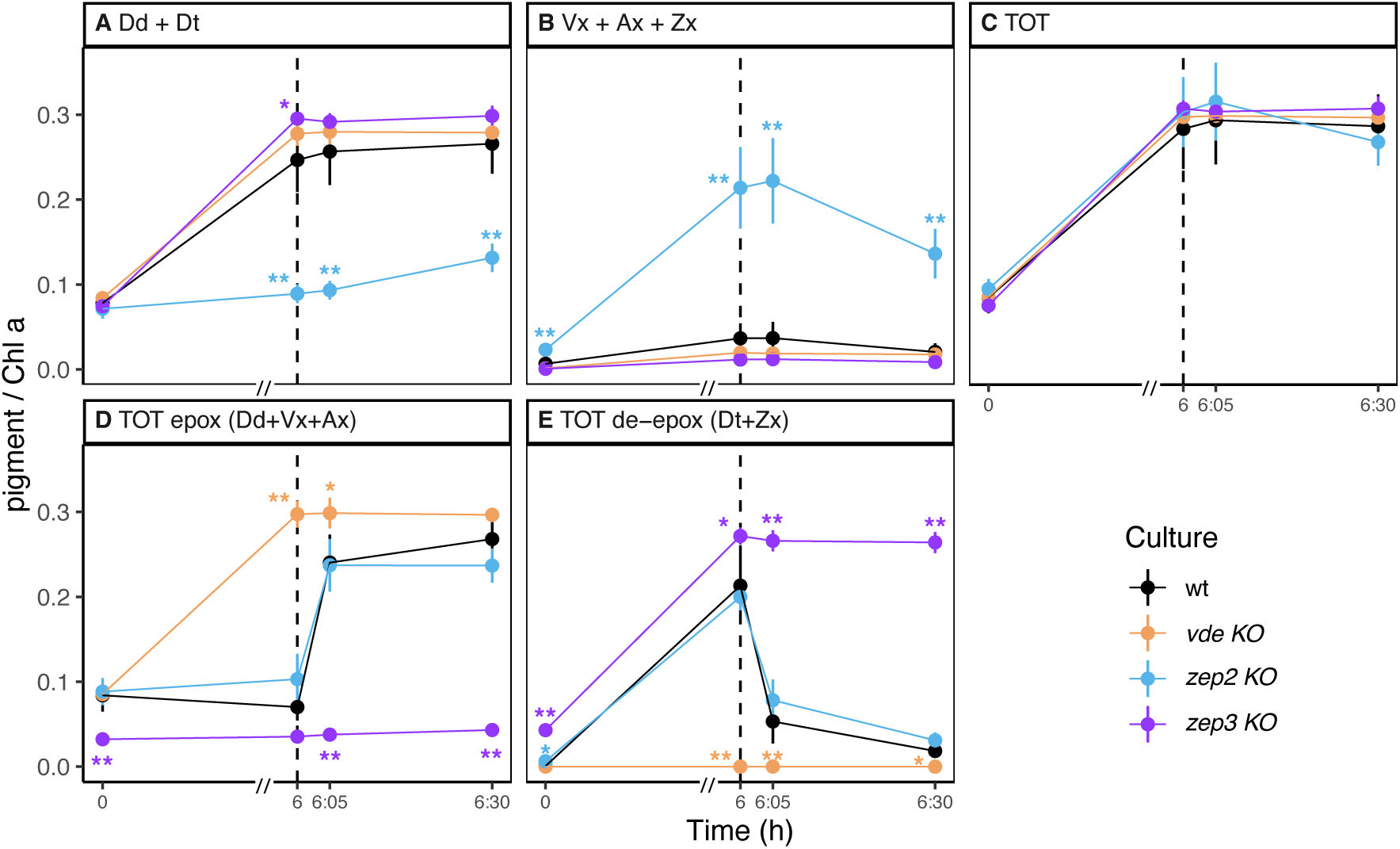
Total pools of Dd and Vx cycle pigments in wt, *vde KO*, *zep2 KO* and *zep3 KO* after 5 days of HL:LL regime (average ± sd, n=3). Pigment content is expressed as pigment:chlorophyll *a* ratio (mol/mol). Time 0 is defined as the beginning of the HL phase (i.e. the sampling point in LL that immediately precedes the onset of HL on the fifth day of HL:LL treatment). The 6 h of HL phase and 30 min of recovery phase (LL) are separated with a dashed line. On the x-axis, recovery phase (from 6 to 6:30 h) was artificially enlarged to allow better visualization; axis break is indicated by a double dash (//). (*A*) total pool of Dd cycle pigments (diadinoxanthin + diatoxanthin); (*B*) total pool of Vx cycle pigments (violaxanthin + antheraxanthin + zeaxanthin); (*C*) sum of all Dd and Vx cycle pigments; (*D*) total pool of epoxidized pigments across different xanthophyll cycles (diadinoxanthin + violaxanthin + antheraxanthin); (*E*) total pool of de-epoxidized pigments across different xanthophyll cycles (diatoxanthin + zeaxanthin). Statistical significance marks indicate significant differences between the corresponding mutant line and wt at each time point, according to adjusted p-value of multiple comparison t-test (*: p < 0.05; **: p < 0.005).

Previous studies suggested that *de novo* synthesis from β-carotene (rather than the catabolism of other major carotenoids) is responsible for the high light-induced accumulation of Zx in *P. tricornutum* (Lohr and Wilhelm, 1999) as in other algae (e.g., *Chlamydomonas reinhardtii* (Baroli et al., 2003)). Accordingly, we observed no significant differences in fucoxanthin content under different light conditions between all investigated mutants and wt (**Fig. S7**), thus confirming that *de novo* synthesis supplements xanthophyll cycle pigments during high light.

To further investigate the function-related accumulation of specific xanthophylls, we calculated the total pool of epoxidized (**Fig. 2D**) and de-epoxidized pigments (**Fig. 2E**). In response to 6 h of HL, *vde KO* accumulated a significantly higher amount of epoxidized xanthophylls, while these were almost absent in *zep3 KO*. Instead, *zep3 KO* accumulated a significantly higher amount of de-epoxidized pigments. Despite the strong differences in the individual amounts of Vx and Dd cycle pigments (**Fig. 1**), *zep2 KO* did not display significant differences in the total pools of epoxidized or de-epoxidized xanthophylls throughout the light treatments compared to wt (**Fig. 2E**).

### Diversification of ZEP2 and ZEP3 uncouples *de novo* xanthophyll biosynthesis from xanthophyll cycling

Previously, ZEP2 had been indicated as main candidate for driving Dt epoxidation, based on the strong accumulation of monoepoxides (Ax and lutein epoxide) in a ZEP-deficient *npq2* mutant of *Arabidopsis thaliana* stably transformed with *ZEP2* from *P. tricornutum* (Eilers et al., 2016). Our results however demonstrate that ZEP3 is the ZEP isoform mainly co-regulating the Dd cycle with VDE in *P. tricornutum*. This is consistent with the clustered organization of *VDE* and *ZEP3* within the genomes of *P. tricornutum* and various other diatoms (Coesel et al., 2008; Frommolt et al., 2008; Bai et al., 2022), with several reports of their correlated expression pattern under light stress (Coesel et al., 2008; Nymark et al., 2009; Kuczynska et al., 2020; Kan et al., 2023). The function of ZEP3 was also recently confirmed in other independent studies (Ware et al., 2024; Græsholt et al., 2024).

Although *zep3 KO* accumulated high amounts of Dt during HL, the mutant was almost devoid of Dt before the onset of HL on the 5^th^ day of light stress regime (**Fig. 1**, Time = 0 h), indicating that Dt epoxidation could still be performed in this mutant on longer time scales. Transient expression of *ZEP2* or *ZEP3* from *P. tricornutum* in the ZEP-deficient *aba2* mutant of *Nicotiana plumbaginifolia* lacking epoxy xanthophylls resulted in almost identical pigment phenotypes accumulating Vx, neoxanthin and lutein epoxide (**Supplemental Fig. S11**), indicating that both isoforms can accept the same xanthophylls as substrate and therefore allow epoxidation of Zx as well as Dt. ZEP2 is thus likely responsible for the long-term re-epoxidation of Dt and Zx in *zep3 KO* following HL exposure, although hardly any epoxidation activity was detectable during the first 30 min of recovery. A major contribution of carotenoid degradation to the scavenging of Dt in the *zep3 KO* during prolonged LL recovery appears unlikely, since carotenoid catabolism is dampened under lower irradiances (Biswal, 1995; Steiger et al., 1999).

These observations suggest that ZEP2 and ZEP3 have overlapping but not completely redundant functions. As we reported similarly broad substrate specificities in tobacco (**Supplemental Fig. S11**), we conclude that this difference is most likely not based on substrate preferences of the two enzymes. We propose that *P. tricornutum* separates photoprotective xanthophyll cycling from *de novo* biosynthesis between ZEP3 and ZEP2 respectively, either through functional specialization and/or physical separation of the two isoforms (**Fig. 3**). A detailed version of these models, illustrating the relationship between predicted enzyme function and observed phenotypes is presented in **Fig. S12**.

**Fig. 3.**
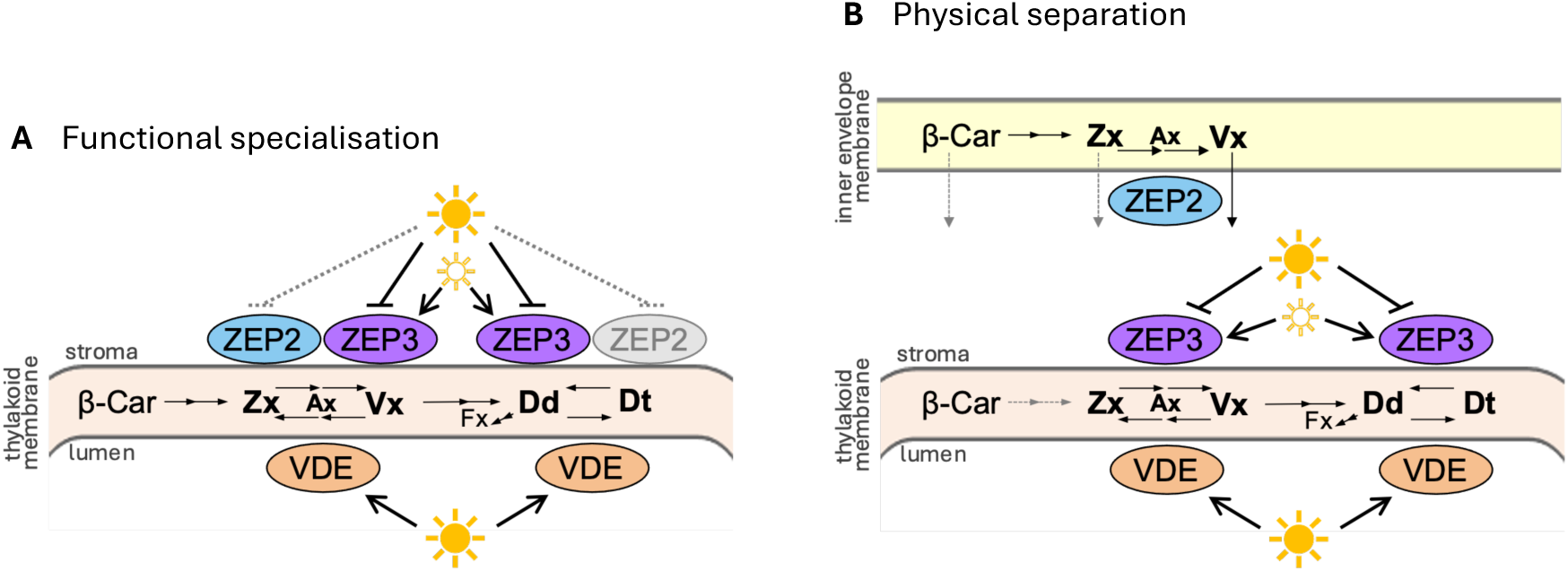
Two models explaining the separation of *de novo* xanthophyll biosynthesis from photoprotective xanthophyll cycling in *P. tricornutum*. In both models, ZEP2 is mostly responsible for *de novo* biosynthesis, while ZEP3 mostly controls photoprotective xanthophyll cycling as antagonist of VDE. (*A*) Functional specialization of ZEP2 and ZEP3. Activity of ZEP3 is maximal under lower irradiances (represented by a hollow sun) and minimal at higher irradiances (represented by a sun), while VDE activity is maximum at higher irradiances. In contrast, ZEP2 is constitutively active, although less efficient than ZEP3 at least for Dt epoxidation. (*B*) In the physical separation model, the two processes are localized in different areas of the chloroplast stroma. Namely, *de novo* biosynthesis of Vx from Zx is catalyzed by ZEP2, confined mostly at the inner envelope, while photoprotective xanthophyll cycling is regulated by ZEP3, located mostly at the thylakoid membrane.

### Despite a decreased Dd+Dt pool, *zep2 KO* displays wt-like NPQ capacity

To investigate the physiological effects of Vx cycle pigments accumulation in *zep2 KO*, the experiment was repeated with an increased number of time points during recovery phase, and pigment analysis was coupled with NPQ measurements. To exclude the influence of the so-called “dark NPQ” (i.e., the formation of NPQ in the dark, typical of diatoms (Goss et al., 2006a; Grouneva et al., 2009)) we used the maximum fluorescence measured in LL (0 h) as Fm for NPQ calculation, which is the common approach in diatoms in contrast to plants as low irradiance represents the condition at which maximum relaxation is achieved (Lepetit et al., 2013).

HPLC analysis confirmed the *zep2 KO* phenotype observed in the first set of experiments (**Fig.1-2**), both in terms of xanthophyll cycle pigments (**Fig. 4A-E**) and total pools (**Supplemental Fig. S9**). The additional time points improved the resolution of the Vx cycle kinetics (**Fig. 4C-E**): while Zx decreased immediately after the cultures were switched from HL to LL, Ax transiently increased and peaked around 10 min of recovery, and then decreased in favor of an increase of Vx.

**Fig. 4.**
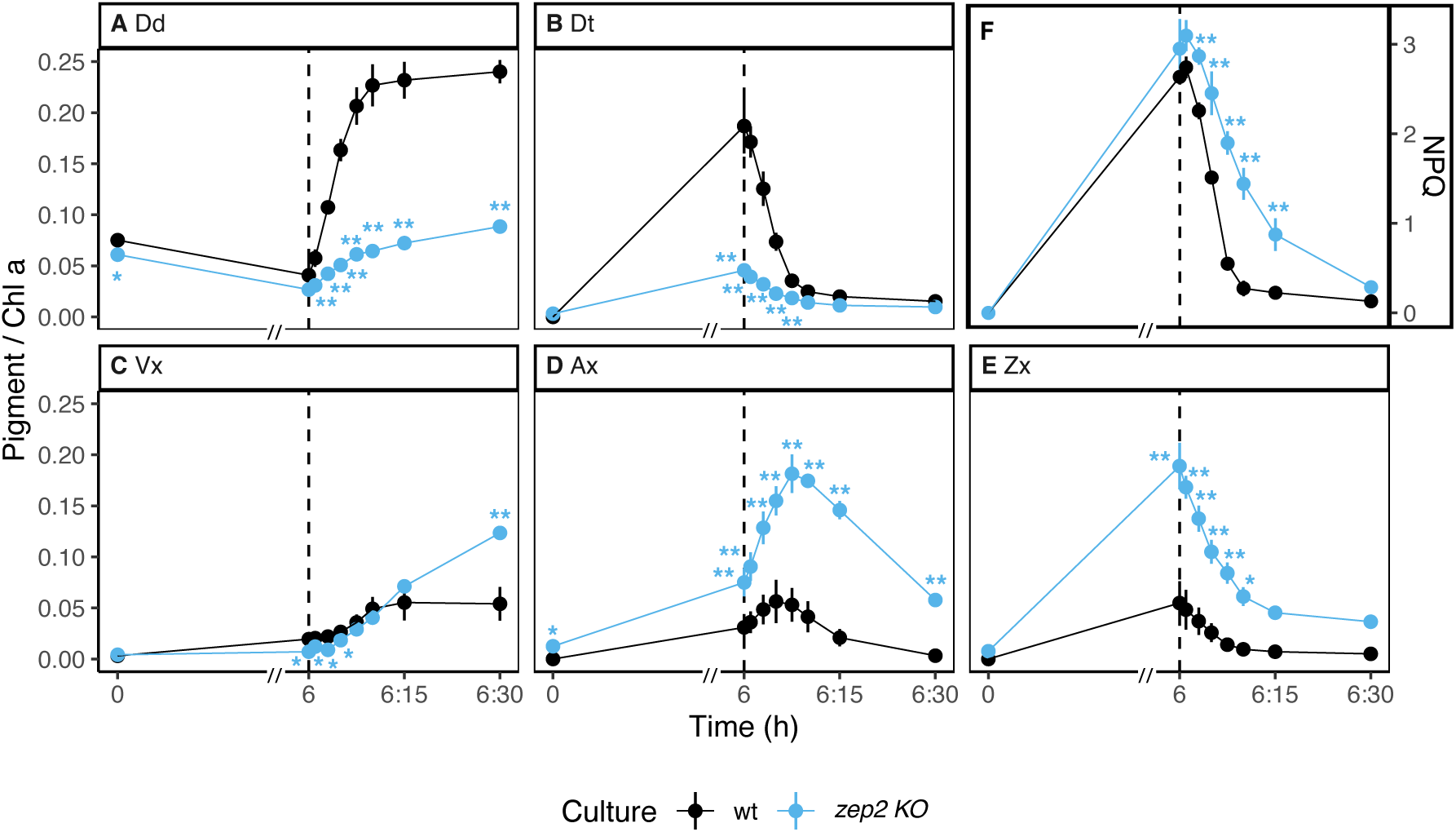
Pigment content of Dd and Vx cycle pigments coupled with NPQ analysis in wt and *zep2 KO* after 5 days of HL:LL regime (average ± sd, n=3). Pigment content is expressed as pigment:chlorophyll *a* ratio (mol/mol). Time 0 is defined as the beginning of the HL phase (i.e. the sampling point in LL that immediately precedes the onset of HL on the fifth day of HL:LL treatment). The 6 h of HL phase and 30 min of recovery phase (LL) are separated with a dashed line. On the x-axis, recovery phase (from 6 to 6:30 h) was artificially enlarged to allow better visualization; axis break is indicated by a double dash (//). (*A*) diadinoxanthin; (*B*) diatoxanthin; (*C*) violaxanthin; (*D*) antheraxanthin; (*E*) zeaxanthin; (*F*) NPQ. Statistical significance marks indicate significant differences between the corresponding mutant line and wt at each time point, according to adjusted p-value of multiple comparison t-test (*: p < 0.05; **: p < 0.005).

Analysis of epoxidation kinetics during LL recovery (**Fig. 5**) showed that Zx and Dt were epoxidized with similar rates within each algal strain, despite *zep2 KO* displayed lower rates for both reactions compared to the wt.

**Fig. 5.**
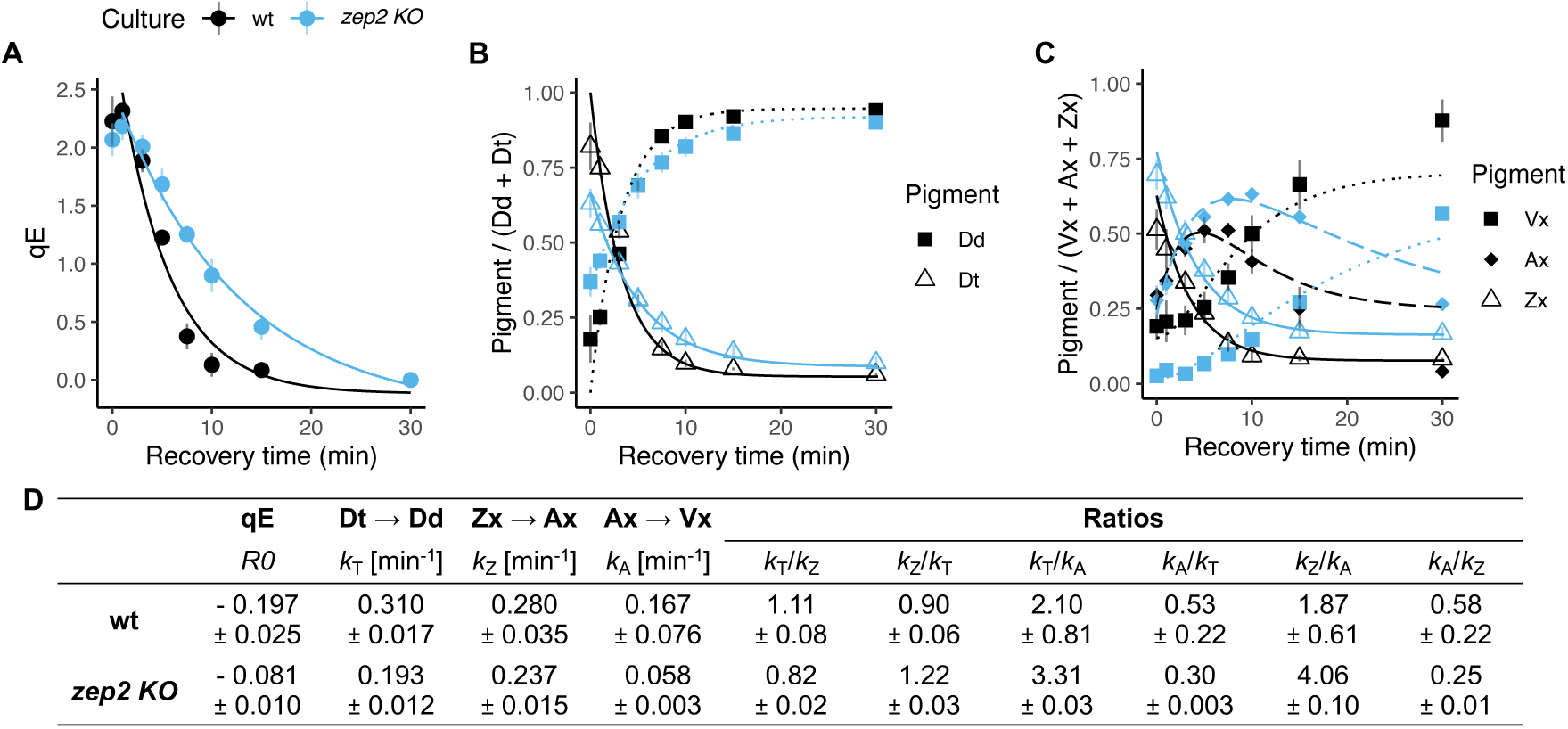
First order kinetics of qE recovery and of re-epoxidation of xanthophyll cycle pigments in wt and *zep2 KO* during 30 min of recovery phase after HL (average ± sd, n=3). (*A*) recovery rates of qE; (*B*) epoxidation of diadinoxanthin (Dd) to diatoxanthin (Dt); (*C*) epoxidation of zeaxanthin (Zx) via antheraxanthin (Ax) to violaxanthin (Vx); (*D*) corresponding rate constants of: qE recovery (R0); Dt to Dd epoxidation (k_T_);, Zx to Ax epoxidation (k_Z_); Ax to Vx epoxidation (k_A_).

Despite a strongly reduced accumulation of Dd cycle pigments during HL (**Fig. 4A-B**), *zep2 KO* still generated the same NPQ as wt (**Fig. 4F**). While NPQ relaxation was initially slower in the mutant, as evidenced by the analysis of qE relaxation kinetics (**Fig. 5A**), no significant differences were observed after 30 min of recovery, indicating that *zep2 KO* was able to relax its NPQ with an efficiency comparable to the wt. These results were confirmed with a second independent *ZEP2* deficient mutant and a wt-like phenotype was restored for *zep2 KO* after complementation with native *ZEP2* (**Supplemental Fig. S9-10**). The same analysis performed on *zep3 KO* showed that the lack of Dt epoxidation during recovery directly corresponds to a lack of NPQ relaxation and a persisting quenching state (**Supplemental Fig. S13-S14**), thus confirming the role of this enzyme as main isoform for the recovery of the Dd cycle.

### The Vx cycle can function as NPQ catalyst in *P. tricornutum*

In most diatoms, qE displays a robust linear correlation to the amount of Dt when no significant amounts of Vx cycle pigments are present (Goss et al., 2006a; Blommaert et al., 2021; Croteau et al., 2021). To factor in the presence of both distinct pools of de-epoxidized xanthophylls, we performed five different calculations of the de-epoxidized pigment pools possibly contributing to NPQ and analyzed their correlation with measured qE (**Fig. 6**).

**Fig. 6.**
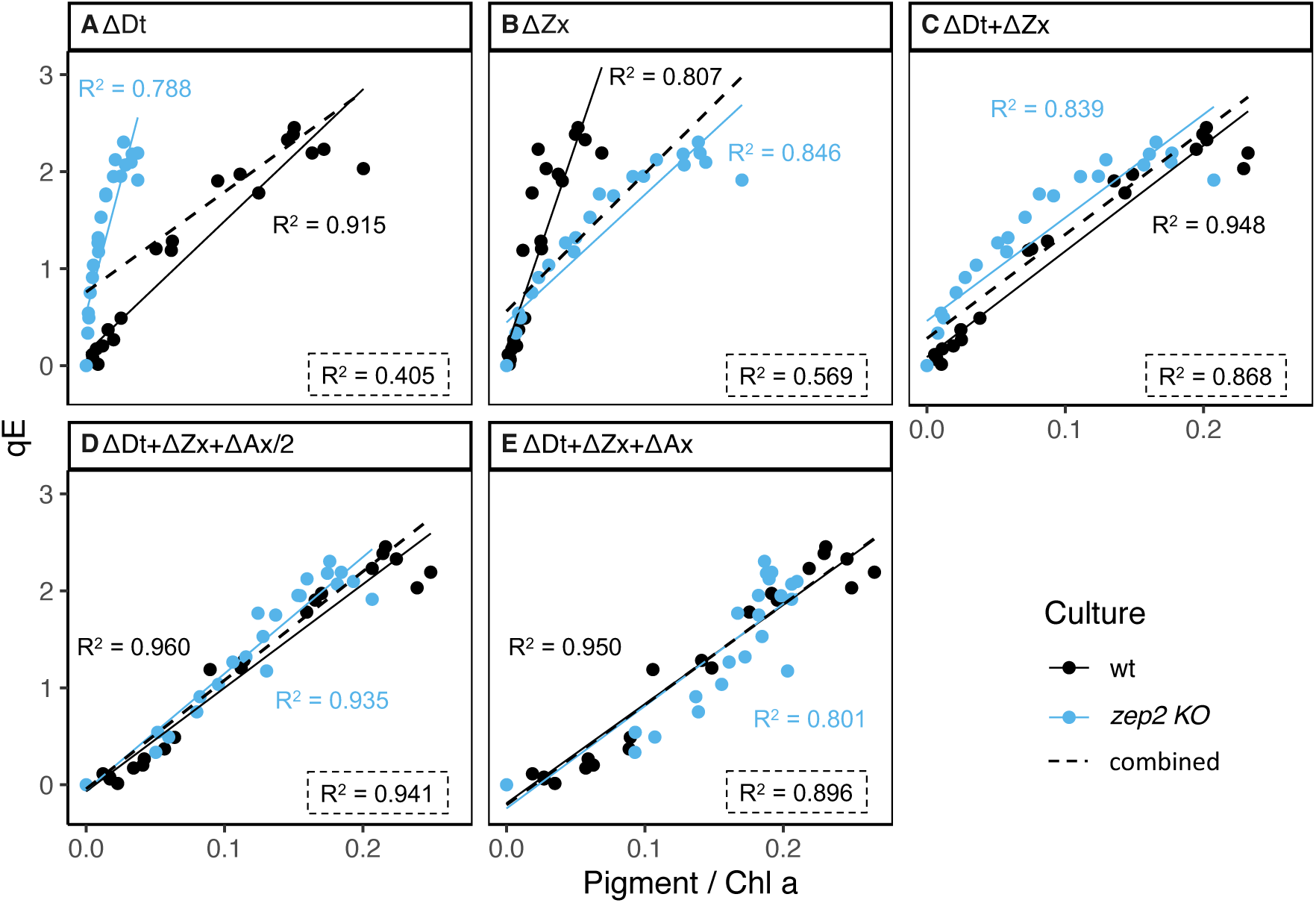
Correlation between qE and de-epoxidized pigments in wt and *zep2 KO* during recovery from 6 h of HL, on the 5^th^ day of HL:LL exposure. Each point represents a separate measurement, repeated for 3 independent biological replicates. For each strain, different calculations of de-epoxidized pigment pool are displayed, with corresponding linear models and R^2^. Dashed lines represent the linear model obtained when considering *zep2 KO* and wt as one unique population (combined); corresponding R^2^ is displayed in the dashed text box at the bottom right corner of each facet. (*A*) diadinoxanthin only; (*B*) zeaxanthin only; (*C*) diatoxanthin and zeaxanthin; (*D*) diatoxanthin, zeaxanthin and ½ antheraxanthin; (*E*) diatoxanthin, zeaxanthin and antheraxanthin.

Dt showed a clear linear relationship with qE in the wt (**Fig. 6A**), while this was much weaker in *zep2 KO*. The opposite was observed with Zx (**Fig. 6B**). In both cases, wt and *zep2 KO* displayed a completely different correlation pattern, noticeable by the poor fit of the combined linear model (i.e., including data points from both wt and *zep2 KO*). With the sum of Dt and Zx (**Fig. 6C**), model fit improved and the slopes of the linear regression for wt and mutant began to converge. The linear fit strongly improved when summing Dt, Zx and ½ Ax (**Fig. 6D**): models for both *zep2 KO* and wt converged towards the general linear model, with the highest R^2^ values. This indicates that the size of this specific pigment pool correlates with qE in a similar manner between these strains. Moreover, the addition of the full Ax pool (**Fig. 6E**) yielded a poorer fit for all investigated lines (with a notable decrease in R^2^ values). We conclude that the sum of Dt, Zx and ½ Ax most adequately describes the xanthophyll cycle-dependent component of NPQ in HL-acclimated *P. tricornutum*. We further confirmed these results with a second *ZEP2* deficient mutant and complementation of *zep2 KO* with native *ZEP2* (**Supplemental Fig. S15**).

## Discussion

In this study, we discovered that pigments of the Vx cycle can contribute to NPQ induction and regulation in algae that normally rely on the Dd cycle, and report that both Zx and Ax can induce NPQ, with the latter having only around half of the quenching-induction capacity of Zx and Dt. Ax participates in the regulation of NPQ in various other photosynthetic eukaryotes, including plants (Gilmore and Yamamoto, 1993; Gilmore et al., 1994; Demmig-Adams and Adams, 1996; Zaks et al., 2013) and green lineage algae (e.g. *Chlorella vulgaris* (Goss et al., 2006b), *Mantoniella squamata* (Goss et al., 1998)). In the heterokont *Nannochloropsis oceanica*, Ax contributes to the onset of qE with about 30% efficiency compared to Zx (Short et al., 2023). Moreover, in the giant kelp *Macrocystis pyrifera*, Ax triggers NPQ with half of the efficiency observed for Zx (Ocampo-Alvarez et al., 2013). Brown algae only possess the Vx cycle (García-Mendoza and Colombo-Pallotta, 2007) but display diatom-like qE (i.e., relying directly on the xanthophyll cycle while not being activated by ΔpH alone (Lavaud and Goss, 2014)), likely explaining the similarities with our findings for *P. tricornutum*.

Xanthophylls are known to compete for different binding sites within the antennae (Croce et al., 1999; Hobe et al., 2000; Caffarri et al., 2001; Mascoli et al., 2021), and mutants lacking one pigment can substitute it with a closely related one (e.g., in *A. thaliana* (Kalituho et al., 2007; Dall’Osto et al., 2007) and *C. reinhardtii* (Azadi-Chegeni et al., 2022)). Vx and Dd cycle pigments are highly similar molecules, that differ only due to a triple bond within the isoprenoid backbone of Dd and Dt, that renders their central polyene moiety asymmetrical (**Fig. 1**). As it is established that Dd and Dt must be bound to light harvesting proteins to regulate NPQ (Schumann et al., 2007; Giovagnetti et al., 2022), we propose that Vx, Ax and Zx, when accumulated in favorable concentrations, could occupy the same binding sites thus contributing to qE induction, as indicated by our data (**Fig. 4-6**). Structural differences between the pigments (including the intrinsic asymmetry of Dd and Dt opposed to the symmetric Vx cycle counterparts) could play a role in the differences in quenching efficiency reported between Ax and Dt/Zx.

These results suggest that Dt might be equally able to trigger NPQ as Zx/Ax do in taxa that normally rely on the Vx cycle. Once the yet unknown Dd synthase (Dautermann et al., 2020) is discovered, this hypothesis could be tested by implementing the Dd biosynthesis pathway in the green lineage.

Xanthophyll cycling is thought to represent an innovation of eukaryotic oxygenic phototrophs, and to have evolved in parallel with light harvesting and other photoprotection strategies (Coesel et al., 2008; Garcia-Mendoza et al., 2011). The Dd cycle most probably evolved in an alga that already possessed the Vx cycle, as Vx is an obligate precursor of Dd and fucoxanthin (Lohr and Wilhelm, 1999; Dautermann et al., 2020; Bai et al., 2022) and VDE and ZEP have sufficiently broad substrate specificities for accepting the xanthophylls from both cycles as substrates (Yamamoto and Higashi, 1978; Bojko et al., 2013; Olchawa-Pajor et al., 2019). According to this scenario, both xanthophyll cycles were functional in the same organism during the early evolution of photosynthetic heterokonts.

Extant algae with the Dd cycle still have the potential to accumulate pigments of the Vx cycle (Lohr and Wilhelm, 1999) which, however, is effectively suppressed under most conditions. One explanation for the prevalence of the Dd cycle in these algae could be that Dt is more effective than Zx in triggering NPQ. Our observation that the Vx cycle pigments can efficiently provide NPQ in a modern diatom species that normally relies on the Dd cycle rejects this hypothesis, while it supports the idea of a gradual shift in photoprotection from the Vx to the Dd cycle in heterokont algae.

Replacing the two-step Vx cycle with the one-step Dd cycle could provide the benefit of switching faster between NPQ and photosynthetic light harvesting (Lohr and Wilhelm, 1999; Lavaud et al., 2002; Goss et al., 2006a). In support of this hypothesis, our data show that the kinetics of recovery of qE and of xanthophyll re-epoxidation during the first 15 min of recovery are slower in a culture that accumulates primarily Vx cycle pigments (**Fig. 5**). Moreover, the formation of fucoxanthin involves only 5 enzymatic steps from Dt versus 8 steps from Zx (Bai et al., 2022; Cao et al., 2023), thereby also putatively accelerating the conversion of photoprotective xanthophylls accumulated under higher irradiance into light harvesting pigments when light becomes limited. We postulate that a selective advantage conferred by these factors could have contributed to the success of diatoms in dynamic ecosystem such as turbulent waters that are characterized by frequent and extreme light changes (Lavaud, 2007).

## Materials and Methods

### CRISPR/Cas9-based gene knockout

*Phaeodactylum tricornutum* (Pt1; CCAP1055) maintained for several years in our laboratory was used as reference wt and for the generation of all mutant strains. Independent knockout lines (two for *ZEP2* [Phatr2_56488; chr_1:2222646-2224755(+)], and one for *VDE* [Phatr2_44635; chr_4:1116193-1117626(-)] and *ZEP3* [Phatr2_56492; chr_4:1118248-1120587(+)]) were obtained with CRISPR/Cas9 genome editing, through bacterial conjugation. sgRNAs were designed with CRISPOR (Concordet and Haeussler, 2018) and episomal vectors based on pPtPuc3m diaCas9_sgRNA including the respective sgRNA templates were created following previously published methods (Nymark et al., 2017; Sharma et al., 2018). sgRNA coding sequences can be found in **Supplemental Table S3**. Final episomes (containing CRISPR/Cas9 machinery and zeocin resistance cassette) were first transformed into *E. coli* DH10β cells carrying the mobilization helper plasmid pTA-Mob and further introduced into wtPt1 *via* bacterial conjugation, as previously described (Karas et al., 2015; Diner et al., 2016; Sharma et al., 2018).

Transformed clones appeared 2-3 weeks after conjugation on agar selection plates (16 g/L salinity f/2 + 100 µg/mL zeocin) incubated at 20°C and 100 µmol photons m^-2^ s^-1^ of white light and were transferred on fresh selection plates (16 g/L salinity f/2 agar + 75 µg/mL zeocin). Screening of mutants followed the strategy previously described (Nymark et al., 2017) and included pre-screening of potentially mutated DNA sequences by PCR amplification via DreamTaq (ThermoFisher Scientific Inc., Germany) and high-resolution melting curve analysis (HRM) using Biozym HRM PCR Mix (Biozym Scientific GmbH, Germany); primers are listed in **Supplemental Table S1**. NPQ phenotypes of the putative mutants were also screened through a chlorophyll *a* fluorescence induction-relaxation protocol (5-10 min light stress at 500-600 µmol photons m^-2^ s^-1^ with blue light, followed by recovery at 28 µmol photons m^-2^ s^-1^) performed with an IMAGING-PAM fluorometer (Heinz Walz GmbH, Germany, described below). For this purpose, cultures were inoculated in well plates with liquid media (16 g/L salinity f/2 + 50 µg/mL zeocin) and grown for 3-4 days at 100 µmol photons m^-2^ s^-1^ of white light.

After identification of positive candidates, the isolation and screening procedure was repeated 1-3 times to obtain pure monoclonal cultures. We confirmed the mutation in both alleles and the monoclonality of the colonies using TOPO-TA cloning (ThermoFisher Scientific Inc., Germany) followed by sequencing of the individual alleles. Mutated clones with a KO indel on both alleles were selected for experiments. Additional information is available in **Supplemental Note S1**.

KO mutants were maintained for several months at 16 °C and 25 µmol photons m^-2^ s^-1^ of white light with 16:8 h light:dark photoperiod on 16 g/L salinity f/2 agar plates without antibiotic selection, to facilitate the expulsion of the CRISPR/Cas9 episome. In addition, prolonged cultivation of transgenic lines was necessary to achieve a stable phenotype before performing experiments, as it is known that in *P. tricornutum* the effects of mutations can be alleviated via compensatory mechanisms such as upregulation of other genes (Nymark et al., 2021). All KO strains generated for this study were later cryopreserved according to (Sprecher et al., 2023).

### Complementation of mutant lines

For each targeted gene (*VDE*, *ZEP2*, and *ZEP3*) we created a modified pPTbsr vector containing the whole length wt gene (amplified from gDNA) with predicted endogenous promoter and terminator to achieve wt-like gene expression. To avoid targeting of the newly inserted sequence, synonymous mutations were introduced at the site of the protospacer adjacent motif in *ZEP2* and *ZEP3*, while a selected episome-free *vde* mutant strain was used for complementation with *VDE*. Biolistic transformation was performed according to (Kroth, 2007). Transformants were grown on selection plates (8 g/L salinity f/2 agar + 4 µg/mL blasticidin-S) for 1-3 weeks at 20°C with 100 µmol photons m^-2^ s^-1^ of white light. DNA of selected clones was extracted as described in (Nymark et al., 2017) and PCR was performed with ALLin™ Mega HiFi DNA Polymerase (highQu GmbH, Germany). PCR products were sequenced (Microsynth, Balgach, Switzerland) and clones showing the insertion of the complete target gene were selected. Primers are listed in **Supplemental Table S4**. Complementation lines were then cryopreserved according to (Sprecher et al., 2023).

### PCR genotyping

In absence of a commercial antibody for our target proteins, PCR genotyping with high discrimination polymerase (HiDi, MyPols, Germany) was performed to confirm the purity of all selected lines prior experiments. gDNA was extracted with the same protocol described for transformants screening (Nymark et al., 2017) and amplification was performed according to manufacturer’s instructions (with addition of 1 mM MgCl_2_), using allele-discriminating primers flanking the targeting loci and binding specifically in each line (KO, wt and complemented). Two different primer pairs were used to confirm the presence of both mutated alleles (allele 1 and allele 2) in the KO lines. Primers are listed in **Supplemental Table S1**. PCR products were run on 1% or 2% TAE-agarose gels with 1000 bp or 50 bp GeneRuler (ThermoFisher Scientific Inc., Germany) depending on product size. All gels were stained with ROTI GelStain (Roth Gmbh, Germany). Additional information is available in **Supplemental Note S1**.

### Culture conditions

All experimental strains were maintained in a temperature-controlled growth chamber at 18°C in shaking batch cultures (120-150 rpm). Standard f/2 medium (Guillard, 1975) was prepared with 16 g/L Tropic Marine CLASSIC sea salt (Dr. Biener GmbH, Germany) adjusted to 7.5 pH prior autoclaving. Continuous low light (LL, 30-50 µmol_photons_ m^-2^ s^-1^), measured in culture flask filled with f/2 medium with a ULM-500 light meter with spherical sensor (Heinz Walz GmbH, Germany), was supplied with L 18W/954 LUMILUX® DELUXE Daylight tubes (OSRAM GmbH, Germany).

About one week prior to each experiment, cultures were acclimated to semi-chemostatic conditions at 18 °C and continuous LL. For the whole experiment, cultures were maintained at 1-2*10^6^ cells/mL in semi-chemostat by daily dilution with fresh medium based on cell counts (Multisizer 4e coulter counter, BECKMAN COULTER, USA).

### Experimental setup

Semi-chemostat cultures acclimated to LL were treated for 5 consecutive days with 6 h of high light (HL, 1000-1200 µmol photons m^-2^ s^-1^ measured in f/2 medium, corresponding to ∼700 µmol photons m^-2^ s^-1^ in air as used by (Lohr and Wilhelm, 1999), supplied by HO 80W/865 LUMILUX® Cool Daylight, OSRAM GmbH, Germany in addition to the LL lamps), followed by 18 h of LL. For pigments analysis, samples were harvested before (LL control) and after 5 days of HL:LL treatment. At the fifth day, cultures were sampled before the onset of the 6 h HL phase (0 h), after 6 h of HL and during recovery in LL (5 and 30 min). Samples (5 mL) were vacuum filtered (ISOPORE^TM^ 1.2 µm PC membrane filters, 25 mm), flash-frozen in liquid nitrogen and stored at -80 °C for later analysis with HPLC.

To investigate the correlation between the kinetics of xanthophyll cycling and NPQ, the experiment was repeated on *zep* mutants and wt and pigment analysis was coupled with fluorescence measurements. At the fifth day of HL:LL treatment, chlorophyll fluorescence was measured, using an IMAGING-PAM fluorometer (IMAG-S module equipped with IMAG-L LED-Ring-Array and IMAG-K CCD-camera, connected to an IMAG-C control unit, Heinz Walz GmbH, Germany), immediately before the 6 h HL phase (0 h), after 6 h of HL and in the following LL phase after 1, 3, 5, 7.5, 10, 15 and 30 min. The fluorometer was positioned under the light setup (i.e., lamps supplying HL or LL as described above, arranged around the sampling area) without covers, so that the whole sampling and measuring procedure could be performed under the same illumination. Samples (3 mL) were collected from the shaking Erlenmeyer flasks and immediately transferred into a 24-well plate that was positioned below the CCD-camera of the fluorometer, while still receiving the same experimental light (i.e., HL or LL as defined above) during the measurements. Blue actinic illumination from the IMAGING-PAM was never supplied to the cultures, while a weak blue pulse-modulated measuring light (low repetition rate, ca. 1 Hz) was applied to measure chlorophyll fluorescence without altering the photophysiological state of the samples. A saturating pulse of blue light from the IMAGING-PAM (max intensity: 3000 µmol photons m^-2^ s^-1^; width: 800 ms) was supplied to measure maximum fluorescence at each time point described above. Simultaneously, samples (5 mL) were collected from the same Erlenmeyer flasks for pigment analysis and preserved as described above. This approach allowed to determine a precise pigment-NPQ correlation under our experimental conditions. NPQ (Fm-Fm′/Fm′) was derived defining Fm as the maximum fluorescence obtained at the first LL pulse (0 h) (Murchie and Lawson, 2013). Instead of darkness, we considered LL as condition in which the maximum of reaction centers are opened, since in diatoms NPQ is not relaxed in the dark (Lepetit et al., 2013). To isolate its effect from the photoinhibition-related quenching (qI), qE (i.e., the rapidly relaxing component of NPQ, also indicated as qZ in diatoms) was calculated as Fm’’-Fm’/Fm’, where Fm’’ corresponds to the maximum fluorescence at the end of the recovery phase (30 min) (Blommaert et al., 2021). Similarly, corresponding deltas for xanthophyll cycle pigments were calculated as Δp = [p]-[p]’’, where [p] corresponds to the pigment concentration at a given time point, and [p]’’ at the end of the recovery phase (30 min).

### Pigment analysis (HPLC)

Pigments extraction was performed as in (Buck et al., 2019). Samples (80 µL) were separated on a NUCLEOSIL^®^ C18 column (EC 250/4, 300-5, MACHEREY-NAGEL GmbH, Germany) in a LaChrome Elite HPLC equipped with an L-2130 pump module, L-2455 diode array detector, and L-2200 autosampler (VWR International, Germany). Elution was performed at 0.8 mL/min flow rate using a three-phase gradient consisting of eluent A (85% methanol, 15% 0.5M ammonium acetate in water), B (90% acetonitrile, 10% water), and C (100% ethyl acetate) following the protocol established by (Kraay et al., 1992). Classical diatom pigments were identified from known absorption spectra and retention times (Roy et al., 2011) and quantified using the integrated absorption peak area at 440 nm and previously determined conversion factors. Retention times and λ_max_ for each pigment and representative chromatograms for each mutant are reported in **Supplemental Table S2, Fig. S5**. For Vx cycle pigments, preliminary identification was further confirmed via spiking with pure crystalline standards (DHI group, Hørsholm, Denmark). If detected, content of chlorophyllide *a* was always summed to that of chlorophyll *a*. Preliminary data assessment was performed with EZChrome *Elite* (Agilent, USA) and OpenChrome (Lablicate Gmbh, Germany) (Wenig and Odermatt, 2010).

### Transient expression of *P. tricornutum ZEP2* and *ZEP3* in a ZEP-deficient tobacco mutant

Full-length *ZEP2* and *ZEP3* from *P. tricornutum* strain UTEX 646 were PCR-amplified from cDNA using Phusion High-Fidelity DNA Polymerase (Thermo Scientific, Carlsbad, USA) and subcloned into the pGEM-T® Easy Vector (Promega, Mannheim, Germany). PCR-related mutations were excluded by confirming that sequences of the cloned genes were identical to sequencing results from PCR products. For generation of plasmids for *Agrobacterium*-mediated tobacco transformation, the gene fragments encoding the mature proteins were PCR amplified, subcloned into pGEM®-T Easy, excised with AvrII and PacI and ligated into the AvrII and PacI sites of plasmid pPZPbar-tp_AtZEP_-AtZEP (Dautermann and Lohr, 2017), thereby replacing the ZEP gene from *Arabidopsis thaliana* and yielding plasmids pPZPbar-tp_AtZEP_-PtZEP2 and pPZPbar-tp_AtZEP_-PtZEP3. Growth of the ZEP-deficient *aba2* mutant of tobacco *N. plumbaginifolia* (Marin et al., 1996), the *Agrobacterium*-mediated transformation of tobacco leaves with the pPZPbar constructs of *AtZEP*, *PtZEP2* and *PtZEP3*, sampling and pigment analysis of lyophilized tobacco leaf discs was performed as described before (Dautermann and Lohr, 2017). All analyses were done at least in triplicate from independent transformants, with the HPLC chromatograms showing representative results.

### Data analysis and statistics

DNA and protein sequences were analyzed with Geneious 9.1 (Biomatters, New Zealand, 2016). Analysis of pigment and chlorophyll *a* fluorescence data was performed with R 4.2.1 (R foundation for statistical computing, Vienna, Austria). Multiple comparison t-tests were performed to test differences in average pigment content and NPQ between mutant lines and wt at different time points; adjusted p-values (Holm correction for multiple comparisons) were used to evaluate statistical significance. Normality was evaluated prior with a combination of graphical methods (Q-Q plot, residual analysis) and Shapiro-Wilk test. For qE/Δpigment correlations, linear regressions were modeled with a standard linear model function. Model fit was evaluated with graphical methods (Q-Q plot, residual analysis) and R^2^. Finally, xanthophyll cycle and qE recovery kinetics were fitted in Microsoft Excel and Origin respectively, using first order kinetics equations (Laidler, 1970). For xanthophyll cycle rates, at each time point molar ratios of pigment/chlorophyll *a* were normalized to the total pool size of each xanthophyll cycle individually. To exclude the influence of *de novo* biosynthesis processes taking place during the 6 h of HL, we excluded data points corresponding to the beginning of the HL phase (0 h) from qE/Δpigment correlations and from kinetic rates calculations.

## Author Contributions

C.E.G. and B.L. designed the project; C.E.G., M.L. and B.L. designed experiments; A.F.S., M.A.W., and J.M.B. generated constructs for CRISPR-Cas KO and complementation lines; C.E.G., B.L. and M.A.W. generated and screened the mutant lines; C.E.G. performed experiments on *P. tricornutum* and analyzed the data; M.L. calculated and analyzed the xanthophyll cycle’s kinetics. O.D. generated constructs for and performed the experiments on *Nicotiana plumbaginifolia*; C.E.G., M.A.W., P.G.K., M.L., and B.L. concluded the final data interpretation; C.E.G. wrote the initial draft of the manuscript, edited by B.L. and M.L.; The manuscript was revised and completed with contribution of all authors.

## Acknowledgments

The authors acknowledge Benjamin Bailleul for his support in NPQ modelling. We also thank Annette Ramsperger for her contribution to the generation of mutant lines, and Alexander H. Fürst, Selina Pucher and Manuel Müller for their help during preliminary characterization and genotyping of the mutants, in the aim of a practical course. Finally, C.E.G thanks Prof. David Schleheck for the project discussions.

## Data and strains availability statement

Mutant algal strains are cryopreserved in the laboratory of Prof. Peter Kroth (Department of Biology, University of Konstanz, 78457 Konstanz, Germany) and are available upon request. Additional queries can be addressed to the corresponding authors.

