## Supplemental Information for "Both major xanthophyll cycles present in nature can provide Non-Photochemical Quenching in the model diatom *Phaeodactylum tricornutum*"

**Supplemental Note S1. Genetic transformation of *P. tricornutum*.** A CRISPR-Cas9 site directed mutagenesis approach was used to knockout *ZEP2*, *ZEP3* and *VDE*. We applied a combination of high-resolution melting curve (HRM) analysis and Sanger sequencing to identify clones with disrupted native gene(Nymark et al., 2017). Induction-relaxation experiments were also performed in the preliminary screening phases to detect altered NPQ phenotypes. While *VDE* and *ZEP3* deficient lines showed a substantial lack of NPQ induction or recovery respectively, no clear phenotype alterations were observed for *ZEP2* deficient lines (**Fig. S1**). Due to the diploid nature of *P. tricornutum* genome(Bowler et al., 2008), the presence of knockout mutations on both alleles was finally assessed with TOPO cloning (**Fig. S2**). We selected clones that harbored indels of varying lengths resulting in a disrupted gene on both alleles (**Fig. S3**). Following sequential rounds of singularization to obtain monoclonality and promote the loss of the episome, resulting lines were further confirmed with PCR genotyping (**Fig. S4**).

With the site directed mutagenesis approach used in this study(Nymark et al., 2017), random mutations can be inserted in the targeted gene, which often results in small indels(Nymark et al., 2016) not traceable via gel electrophoresis (e.g., **Fig. S2**). Moreover, compared to other approaches (i.e., homology-directed repair (HDR)(Bai et al., 2022) or random genomic integration via biolistic bombardment(Buck et al., 2019)) episomes are transiently maintained and do not stably integrate antibiotic resistance or transgenes into the host genome. This poses a disadvantage during screening but in turn prevents random integrations in the genome and allows the removal of the episome after mutant isolation, to reclaim antibiotic selection markers if required (e.g., for subsequent transformations) and to prevent off-target mutations.

After amplification of the targeted regions with TAQ (**Fig. S2D**), we pre-selected ex-conjugants with high resolution melting curve analysis (HRM) (and chlorophyll *a* fluorescence phenotyping for *vde* and *zep3* mutants). Only a subset of clones showing clear divergence from the wt were further analyzed with TOPO cloning to assess the presence of biallelic mutations. Only frame shifts resulting in premature stop codons or exceptionally large indels were considered as successful KOs. Alignments of final sequencing results of *zep2*, *zep3* and *vde* deficient lines and wt are shown in **Fig. S2A-C**.

We further confirmed these results by PCR genotyping using HiDi polymerase, that selectively discriminates SNPs at the primers' 3′ end. We thus designed primers whose 3′ ends correspond to only one specific sequence (e.g., wt or KO) based on the original TOPO cloning results. We obtained a set of three primer pairs for each target gene: one specific for the native gene (e.g., *VDE*), and one for each allele of the corresponding deficient line (e.g., *vde KO* allele 1 and *vde KO* allele 2). In addition, we designed a primer pair that selectively amplified the newly introduced gene from the complemented line (e.g., *vde C*) based on the sequence of the complementation plasmid (one primer binding within the target gene and one primer binding a sequence of the pPTbsr backbone, absent in the wt). All primer sequences are reported in **Table S1**.

With this approach we were able to reliably discriminate wt from edited alleles in mixed and monoclonal cell populations and thus could proof the absence of the native gene in each of the selected KO lines (**Fig. S4**): native gene amplification (*VDE, ZEP2 and ZEP3*) gave a positive result only for wt DNA, while amplification of both alleles from the mutants was observed only in the corresponding DNA samples. As only exception, the PCR for *zep2 KO* allele 2 showed a positive amplification also in the wt sample (**Fig. S4B)**; however, for this primer pair a size shift (corresponding to the known sequence for this allele) could be observed in *zep2 KO* compared to the wt, while both bands were amplified in the complemented line (*zep2 C*), confirming the genotyping of this allele despite the unpredicted amplification in the wt. Complementation lines were also successfully genotyped: all samples (*vde C*, *zep2 C* and *zep3 C*) showed a positive amplification for both KO alleles and for the primer pairs specific the complemented line. As part of our quality control, PCR genotyping was repeated prior experiments to ensure absence of potential cross-contamination.

Alignments of the protein sequences deduced from *in silico* translation of the edited genes showed that both alleles of each mutant line had disrupted gene products that either resulted from out-of-frame shifts with premature stop codons or from large insertions/deletions (**Fig. S3**). For all target genes we were able to isolate strains with different mutations on the two allele; thus, we did not observe the predominant loss of heterozygosity reported in previous studies(Nymark et al., 2016; Bai et al., 2022). This could be a result of selection or of differences in the transformation and mutagenesis approaches. However, we often observed a rearrangement of the allele-specific SNPs known in our wt type strain, which would be consistent with the high frequency of mitotic interhomolog recombination observed in *P. tricornutum*(Bulankova et al., 2021).

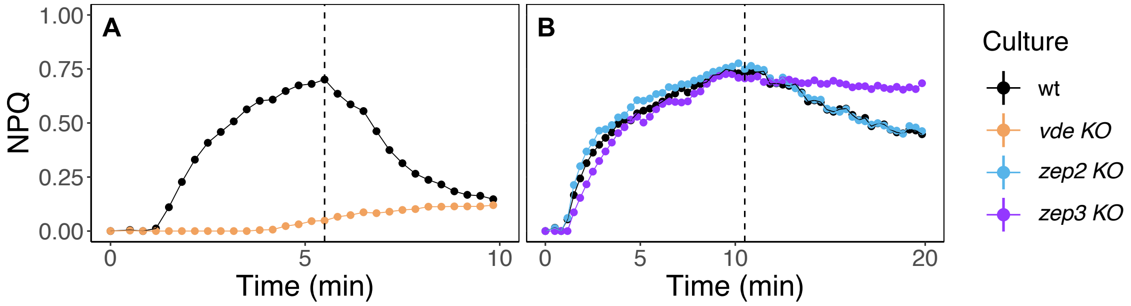


**Fig. S1: Preliminary induction-relaxation screening of *VDE*, *ZEP2* and *ZEP3* deficient lines and wt (representative example).** The first pulse in the dark was followed (after 30 s) by light stress induction and recovery in low light. The switch from light stress to recovery is indicated by a dashed line. (*A*) screening of a *vde* deficient line (500m µmol photons $m^{-2}$ $s^{-1}$ for 5 min, followed by recovery at 28 µmol photons $m^{-2}$ $s^{-1}$); (*B*) screening of *zep2* and *zep3* deficient lines (645m µmol photons $m^{-2}$ $s^{-1}$ for 10 min, followed by recovery at 28 µmol photons $m^{-2}$ $s^{-1}$).

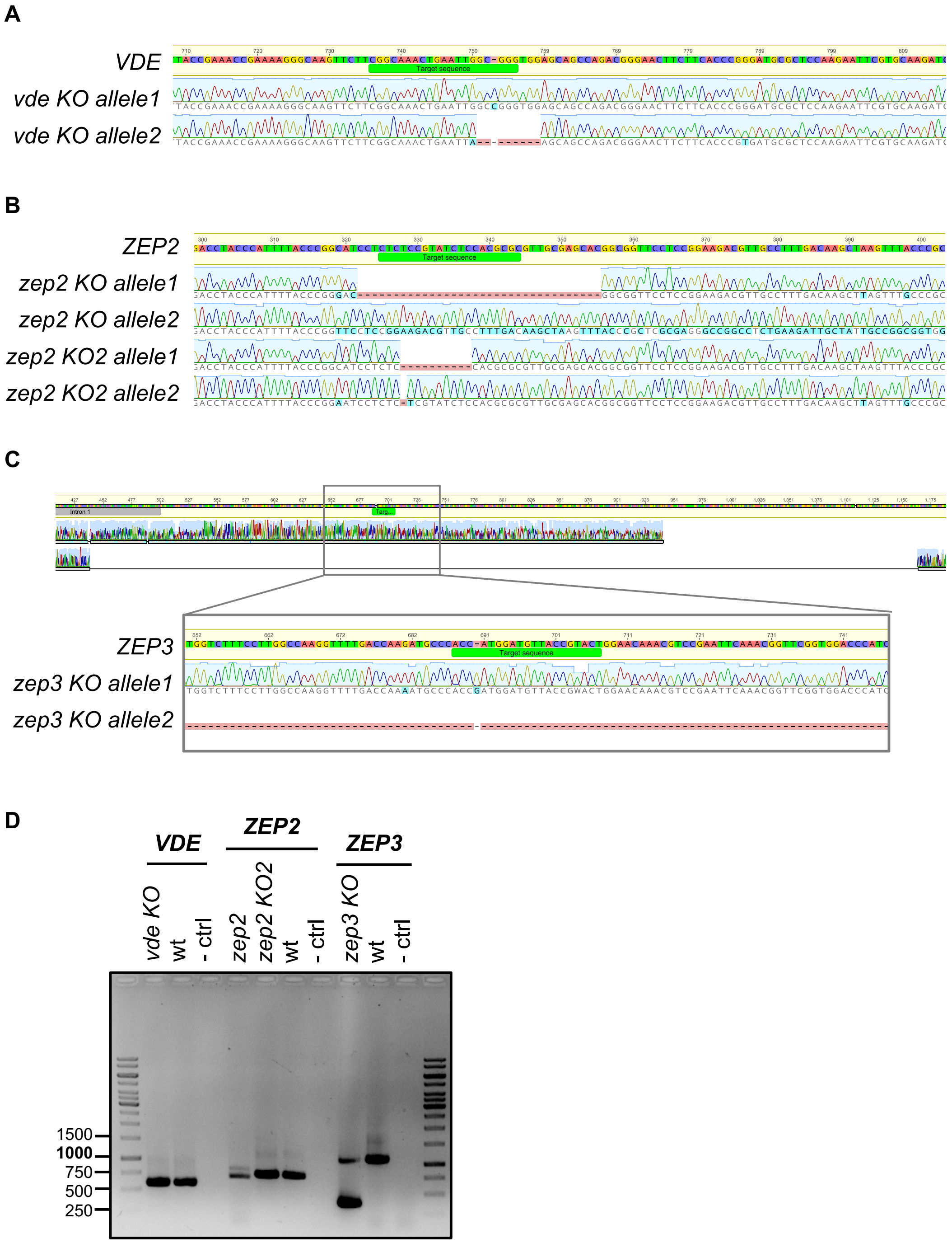


**Fig. S2: Partial DNA sequence alignments of both alleles of the respective target genes for all KO mutants generated in this study.** Alignments were generated with Geneious 9.1 (Biomatters, New Zealand, 2016). Differences from wild type (*Pt*1; CCAP1055) are highlighted. (*A*) *zep2 KO* and *zep2 KO2*; (*B*) *vde KO*; (*C*) *zep3 KO* (*D*) gel electrophoresis of the corresponding TAQ PCR products (1% TAE-agar stained with Roti GelStain) before TOPO cloning.

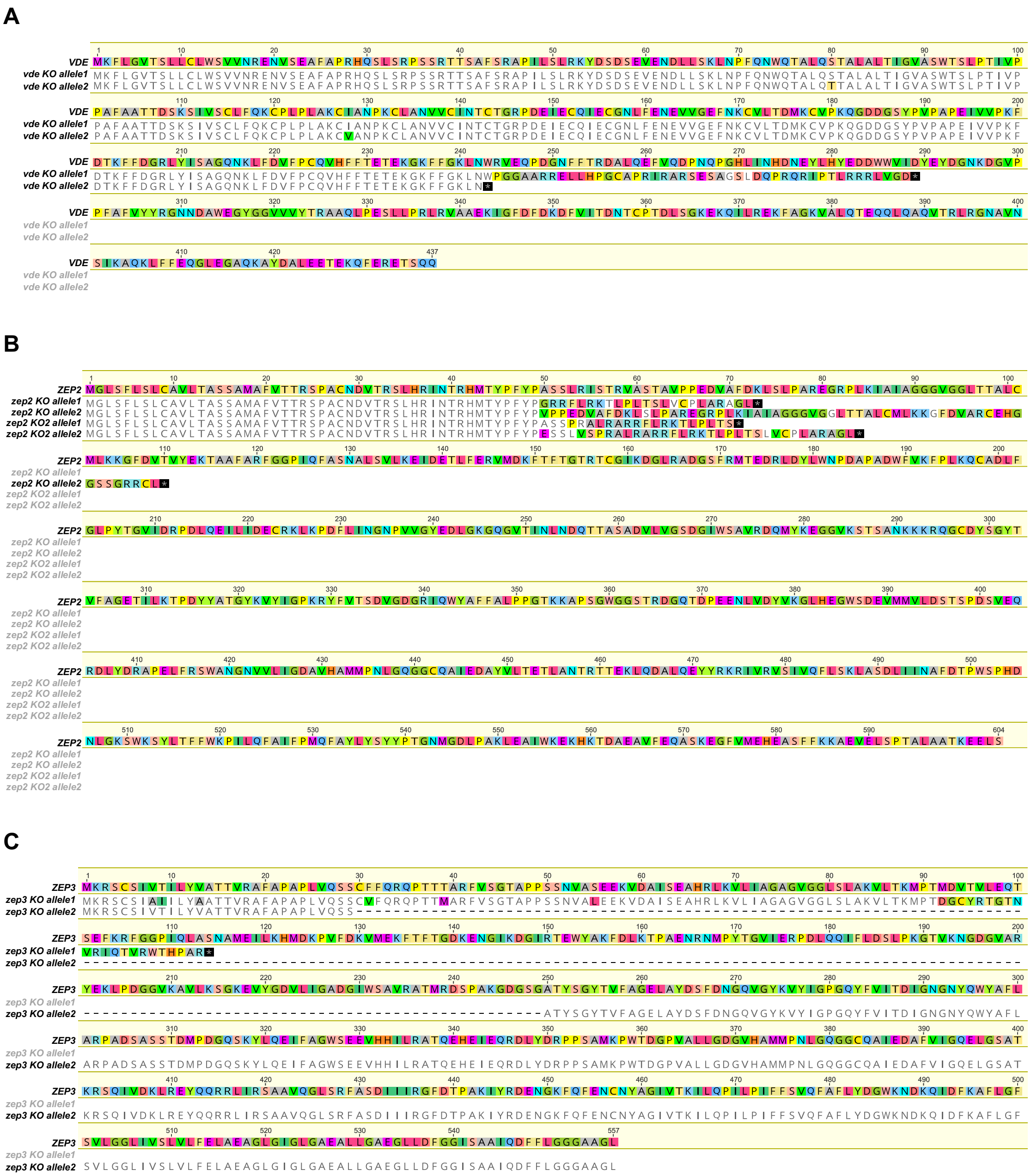


**Fig. S3: Complete protein sequence alignments of both allelic gene products for all knockout mutants generated in this study.** Differences from wild type (*Pt*1; CCAP1055) are highlighted. (*A*) vde KO; (*B*) *zep2 KO* and *zep2 KO2*; (*C*) *zep3 KO*. Alignments were generated with Geneious 9.1 (Biomatters, New Zealand, 2016).

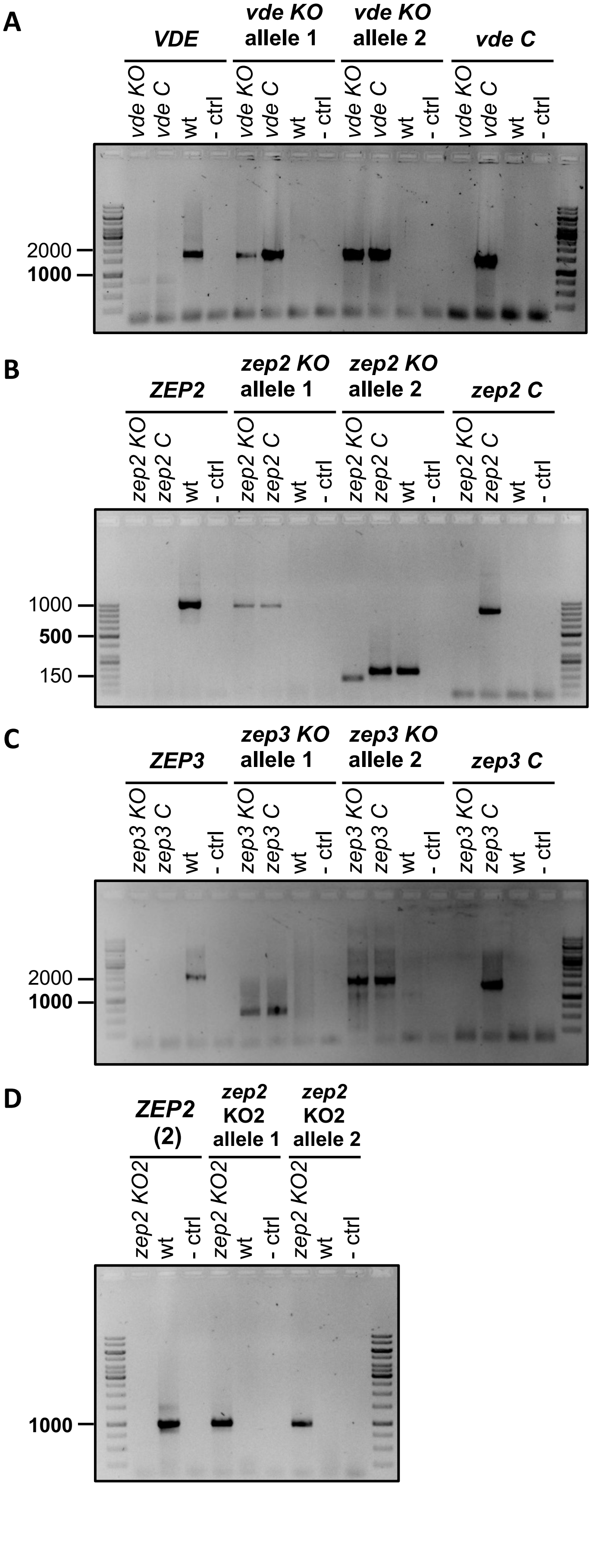
A
VDE
vde KO allele 1
vde KO allele 2
vde C
vde KO
vde C
wt
- ctrl
2000
1000
B
ZEP2
zep2 KO allele 1
zep2 KO allele 2
zep2 C
zep2 KO
zep2 C
wt
- ctrl
1000
500
150
C
ZEP3
zep3 KO allele 1
zep3 KO allele 2
zep3 C
zep3 KO
zep3 C
wt
- ctrl
2000
1000
D
ZEP2 (2)
zep2 KO2 allele 1
zep2 KO2 allele 2
zep2 KO2
wt
- ctrl
1000

**Fig. S4: Molecular characterization of transformed *P. tricornutum* mutants (KO and complementation lines) via PCR genotyping.** Amplification from genomic DNA was performed according to HiDi manufacturer's instructions, using allele-specific primers binding specifically in each line (KO, wt and complemented). For KO lines, two different primer pairs were used to confirm the presence of both mutated alleles (allele 1 and allele 2). Corresponding primer sequences and additional information are available in **Table S1** and **Supplemental Text S1**. For each gel, target sequences are indicated in **bold**, while sample names are placed vertically above each lane. Negative controls (- ctrl) were performed with nuclease-free water instead of DNA. (*A*) amplification of native *VDE*, *vde KO* allele 1, *vde KO* allele 2 and *vde C* specific sequences from *vde KO*, *vde C* and wt DNA samples; (*B*) amplification of native *ZEP2*, *zep2 KO* allele 1, *zep2 KO* allele 2 and *zep2 C* specific sequences from *zep2 KO*, *zep2 C* and wt DNA samples; (*C*) amplification of native *ZEP3*, *zep3 KO* allele 1, *zep3 KO* allele 2 and *zep3 C* specific sequences from *zep2 KO*, *zep3 C* and wt DNA samples; (*D*) amplification of native *ZEP2*, *zep2 KO2* allele 1 and *zep2 KO2* allele 2 specific sequences from *zep2 KO2* and wt DNA samples.

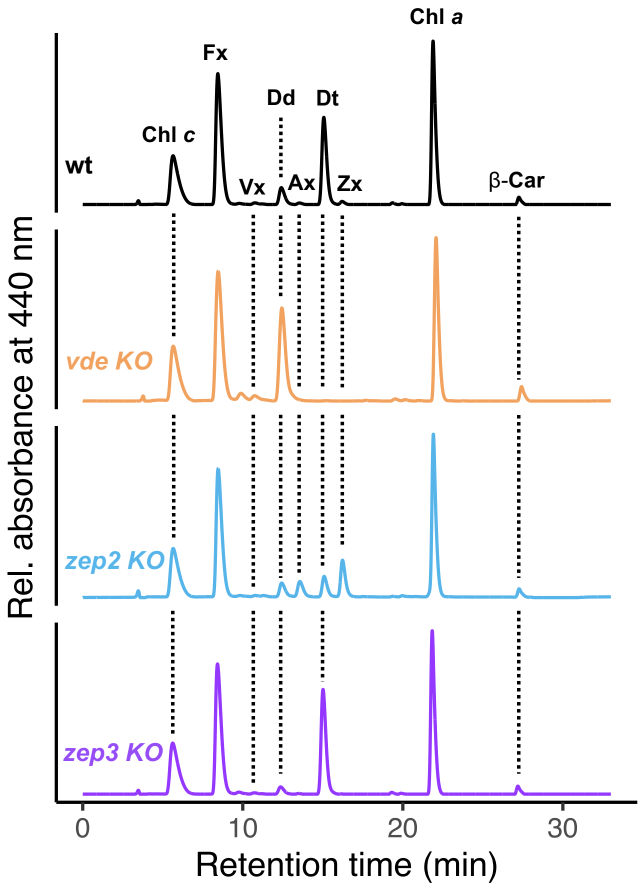


**Fig. S5. Representative chromatograms of pigments extracts from wt, *vde KO*, *zep2 KO* and *zep2 KO* after 6 h of HL.** Chl c: chlorophyll *c*; Fx: fucoxanthin; Vx: violaxanthin; Dd: diadinoxanthin; Ax: antheraxanthin; Dt: diatoxanthin; Zx: zeaxanthin; Chl a: chlorophyll *a*; β-car: β,β-carotene**.**

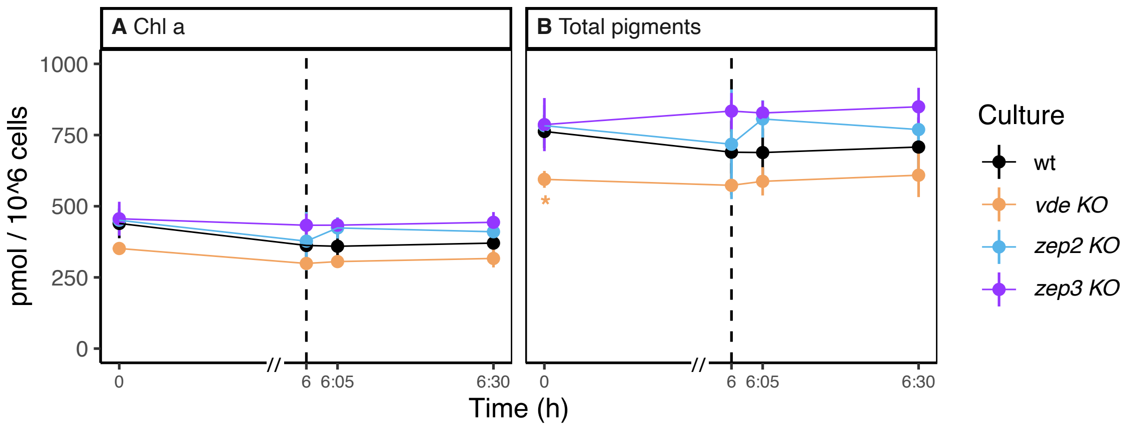


**Fig. S6: Content of chlorophyll *a* and total pigments in wt, *vde KO*, *zep2 KO* and *zep3 KO* after 5 days of HL:LL regime (average ± sd, n=3).** Time 0 is defined as the beginning of the HL phase (i.e. the sampling point in LL that immediately precedes the onset of HL on the fifth day of HL:LL treatment). The 6 h of HL phase and 30 min of recovery phase (LL) are separated with a dashed line. On the x-axis, recovery phase (from 6 to 6:30 h) was artificially enlarged to allow better visualization; axis break is indicated by a double dash (//). (*A*) chlorophyll *a*; (*B*) total pigments. Statistical significance marks indicate significant differences between the corresponding mutant line and wt at each time point, according to adjusted p-value of multiple comparison t-test (*: $p < 0.05$).

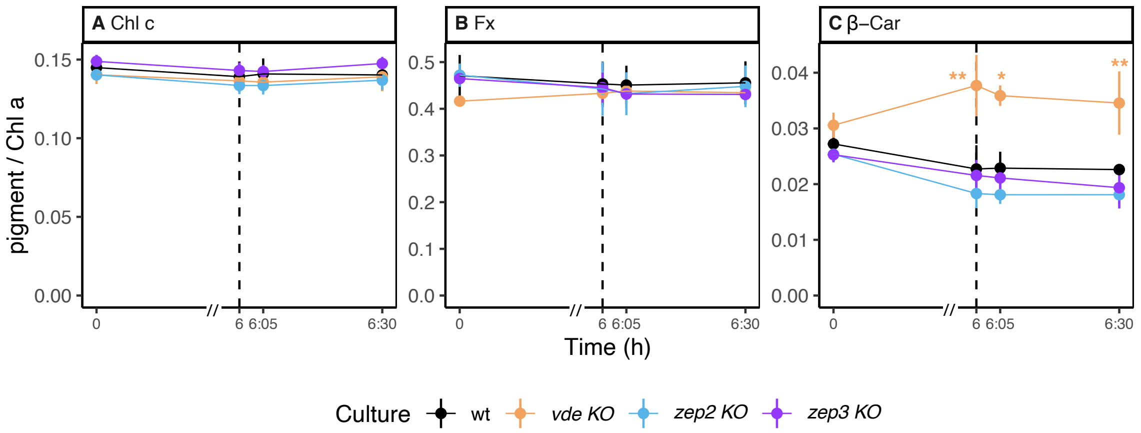


**Fig. S7: Content of other photosynthetic pigments and carotenes in wt, *vde KO*, *zep2 KO* and *zep3 KO* after 5 days of HL:LL regime (average ± sd, n=3).** Pigment content is expressed as pigment per chlorophyll *a* (mol/mol). Time 0 is defined as the beginning of the HL phase (i.e. the sampling point in LL that immediately precedes the onset of HL on the fifth day of HL:LL treatment). The 6 h of HL phase and 30 min of recovery phase (LL) are separated with a dashed line. On the x-axis, recovery phase (from 6 to 6:30 h) was artificially enlarged to allow better visualization; axis break is indicated by a double dash (//). (*A*) chlorophyll *c*; (*B*) fucoxanthin; (*C*) β,β-carotene. Statistical significance marks indicate significant differences between the corresponding mutant line and wt at each time point, according to adjusted p-value of multiple comparison t-test (*: $p < 0.05$ ; **: $p < 0.005$).

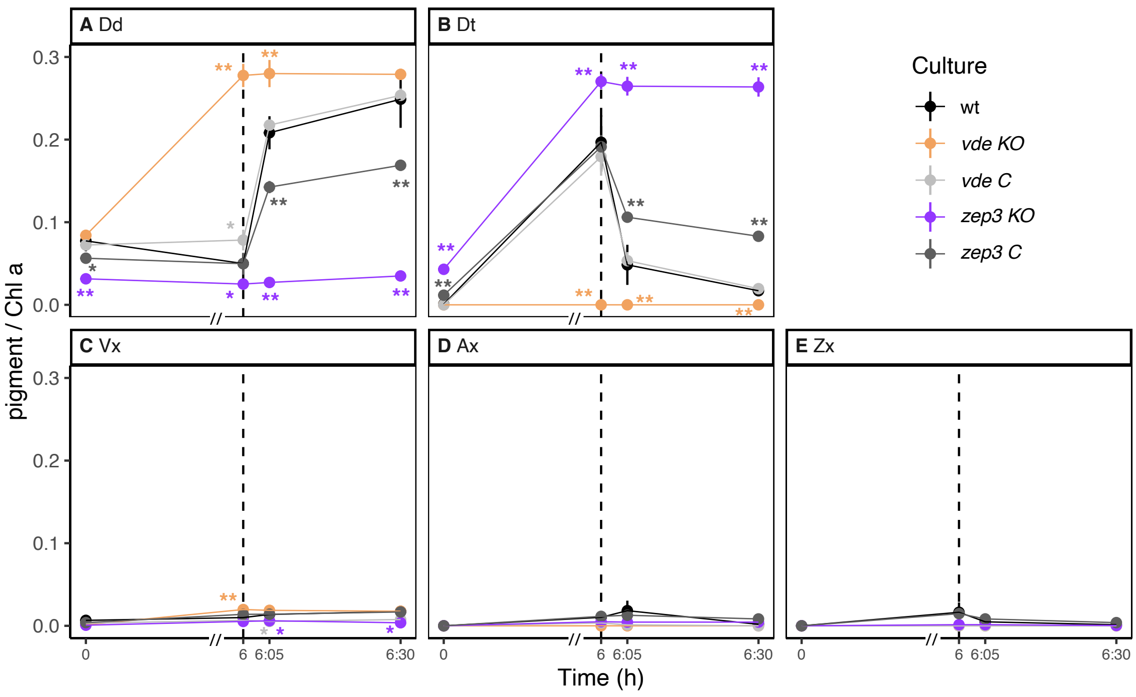


**Fig. S8: Content of Dd and Vx cycle pigments in *vde KO* and *zep3 KO* with respective complementation lines (*vde C*, *zep3 C*) and wt after 5 days of HL:LL regime (average ± sd, n=3).** Pigment content is expressed as pigment per chlorophyll *a* (mol/mol). Time 0 is defined as the beginning of the HL phase (i.e. the sampling point in LL that immediately precedes the onset of HL on the fifth day of HL:LL treatment). The 6 h of HL phase and 30 min of recovery phase (LL) are separated with a dashed line. On the x-axis, recovery phase (from 6 to 6:30 h) was artificially enlarged to allow better visualization; axis break is indicated by a double dash (//). (*A*) diadinoxanthin; (*B*) diatoxanthin; (*C*) violaxanthin; (*D*) antheraxanthin; (*E*) zeaxanthin. Statistical significance marks indicate significant differences between the corresponding mutant line and wt at each time point, according to adjusted p-value of multiple comparison t-test (*: $p < 0.05$ ; **: $p < 0.005$).

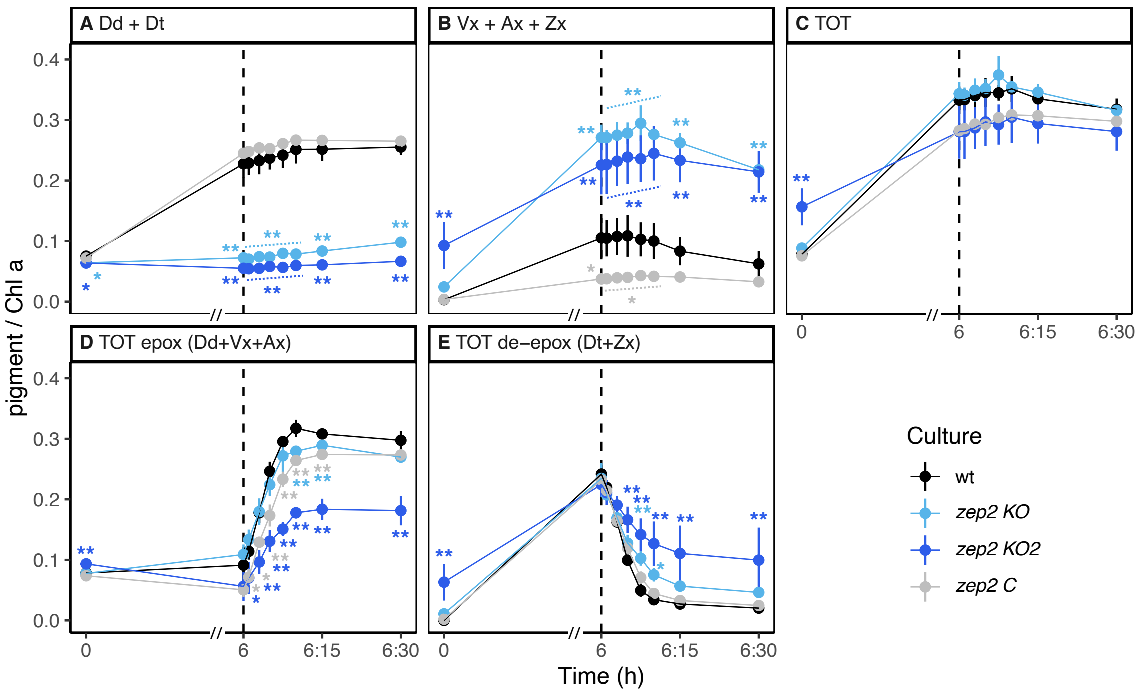


**Fig. S9. Total pools of Dd and Vx cycle pigments in wt, *zep2 KO*, *zep2 KO2* and *zep2* C after 5 days of HL:LL (average ± sd, n=3).** Pigment content is expressed as pigment per chlorophyll *a* (mol/mol). Time 0 is defined as the beginning of the HL phase (i.e. the sampling point in LL that immediately precedes the onset of HL on the fifth day of HL:LL treatment). The 6 h of HL phase and 30 min of recovery phase (LL) are separated with a dashed line. On the x-axis, recovery phase (from 6 to 6:30 h) was artificially enlarged to allow better visualization; axis break is indicated by a double dash (//). (*A*) total pool of Dd cycle pigments (diadinoxanthin + diatoxanthin); (*B*) total pool of Vx cycle pigments (violaxanthin + antheraxanthin + zeaxanthin); (*C*) sum of all diadinoxanthin and Vx cycle pigments; (*D*) total pool of epoxidized pigments across different xanthophyll cycles (diadinoxanthin + violaxanthin + antheraxanthin); (*E*) total pool of de-epoxidized pigments across different xanthophyll cycles (diatoxanthin + zeaxanthin). Statistical significance marks indicate significant differences between the corresponding mutant line and wt at each time point, according to adjusted p-value of multiple comparison t-test (*: $p < 0.05$; **: $p < 0.005$).

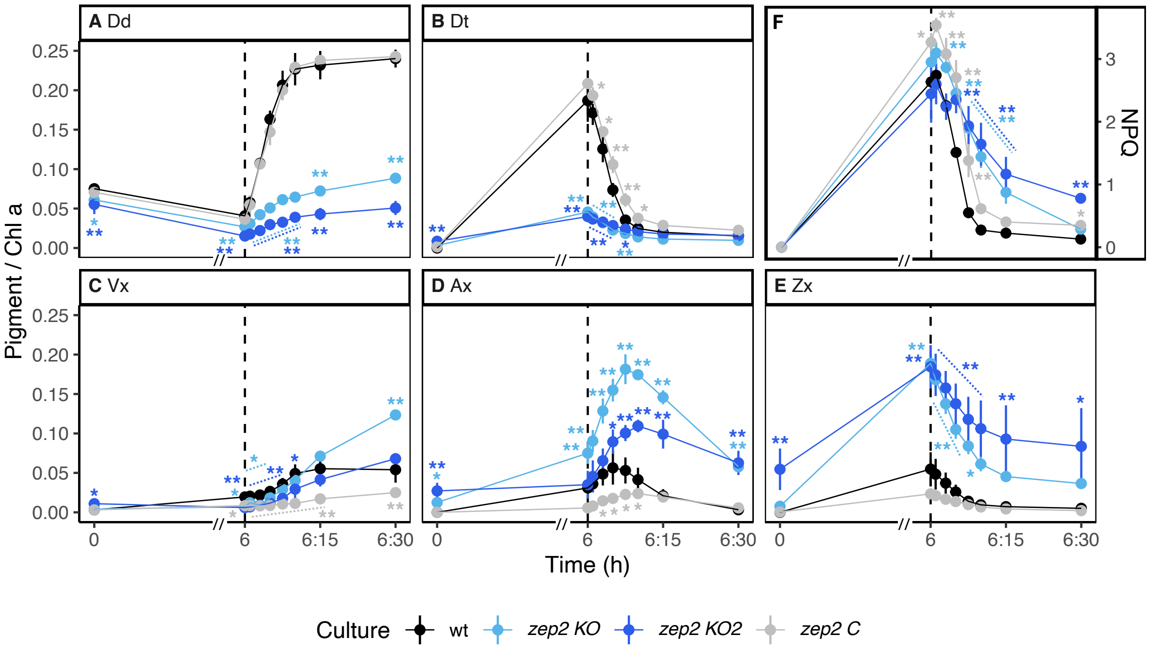


**Fig. S10: Pigment content of Dd and Vx cycle pigments coupled with NPQ analysis in wt, *zep2 KO*, *zep2 KO2* and *zep2 C* after 5 days of HL:LL regime (average ± sd, n=3).** Pigment content is expressed as pigment:chlorophyll *a* ratio (mol/mol). Time 0 is defined as the beginning of the HL phase (i.e. the sampling point in LL that immediately precedes the onset of HL on the fifth day of HL:LL treatment). The 6 h of HL phase and 30 min of recovery phase (LL) are separated with a dashed line. On the x-axis, recovery phase (from 6 to 6:30 h) was artificially enlarged to allow better visualization; axis break is indicated by a double dash (//). (*A*) diadinoxanthin; (*B*) diatoxanthin; (*C*) violaxanthin; (*D*) antheraxanthin; (*E*) zeaxanthin; (*F*) NPQ. Statistical significance marks indicate significant differences between the corresponding mutant line and wt at each time point, according to adjusted p-value of multiple comparison t-test (*: $p < 0.05$; **: $p < 0.005$).

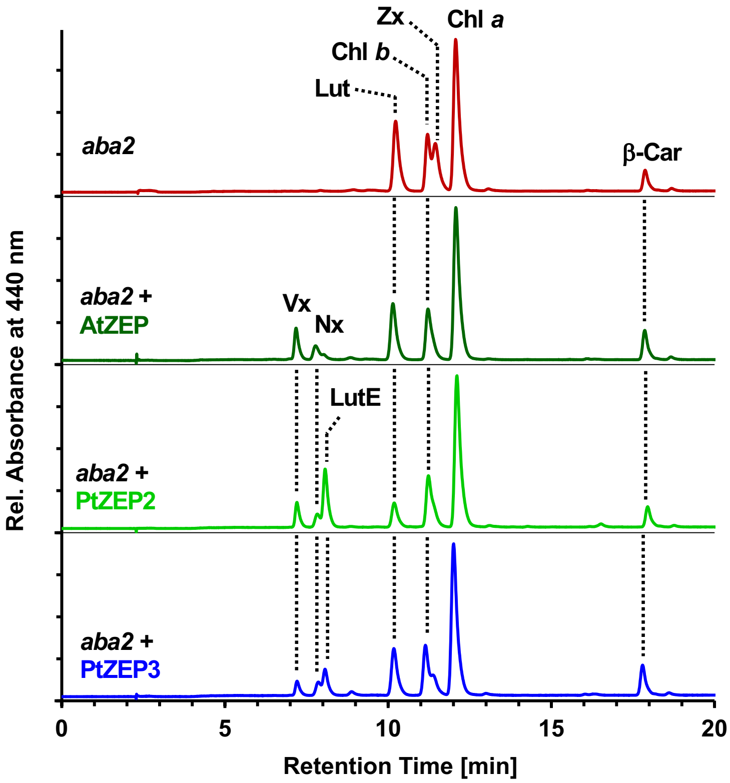


**Fig. S11. Transient expression of *ZEP2* or *ZEP3* from *P. tricornutum* in leaves of the ZEP-deficient *aba2* mutant of tobacco (*Nicotiana plumbaginifolia*) resulted in the formation of violaxanthin (Vx), its derivative neoxanthin (Nx), and lutein epoxide (LutE).** Transient expression of the *ZEP* from Arabidopsis thaliana (AtZEP) as control yielded only Vx and Nx. Other pigments: β-Car, β,β-carotene; Chl, chlorophyll; Lut, lutein; Zx, zeaxanthin.

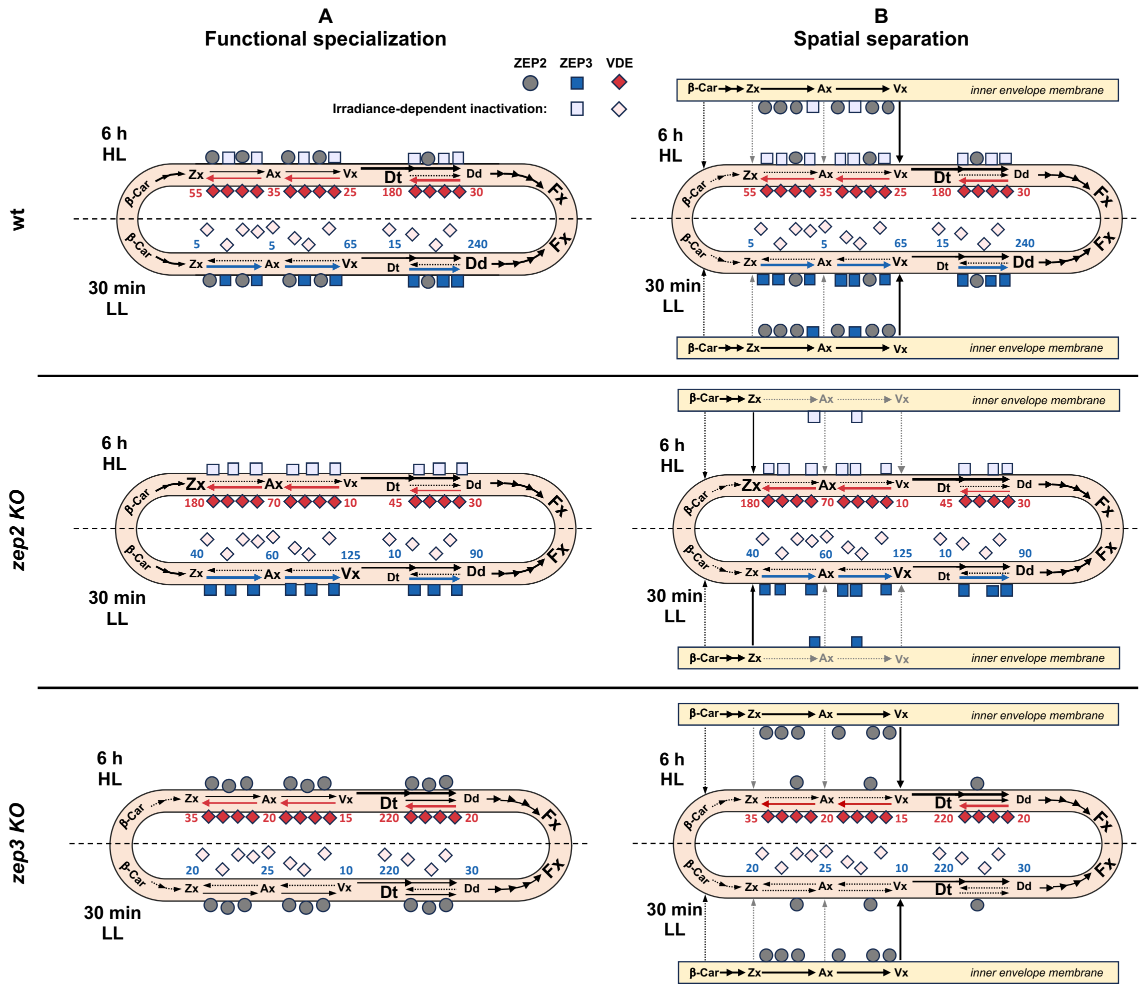


**Fig. S12. Detailed models explaining the observed differences in xanthophyll accumulation between *zep2 KO*, *zep3 KO* and wt.** Different panels show the state of each line at 6h of HL treatment (top) and after 30 min of recovery under LL (bottom). Colored arrows show preferential direction of xanthophyll cycles under HL (red) or LL (blue), numbers represent the relative concentration (xanthophyll:1000 chlorophyll *a*) of each pigment at the corresponding time point. (*A*) Selective accumulation of Vx/Ax/Zx in *zep2 KO* explained by functional differences in the activity of ZEP2 and ZEP3. In this scenario, ZEP3 is selectively inactivated by HL while ZEP2 displays a lower but constitutive activity also under higher irradiances. Lack of ZEP2 causes the accumulation of Zx/Ax in *zep2 KO* under HL, while absence of ZEP3 determines lack of rapid Dt epoxidation in *zep3 KO* during LL recovery. (*B*) The two *zep* phenotypes as result of the different localization of ZEP2 and ZEP3 in the chloroplast. In this model, ZEP2 is mostly located at the inner envelope, where it is responsible for the *de novo* synthesis of Vx from Zx. Therefore, lack of this isoform causes accumulation of Zx. On the contrary, ZEP3 is confined mainly at the thylakoid membranes where it performs xanthophyll cycle-related Dt epoxidation, for which this enzyme is essential.

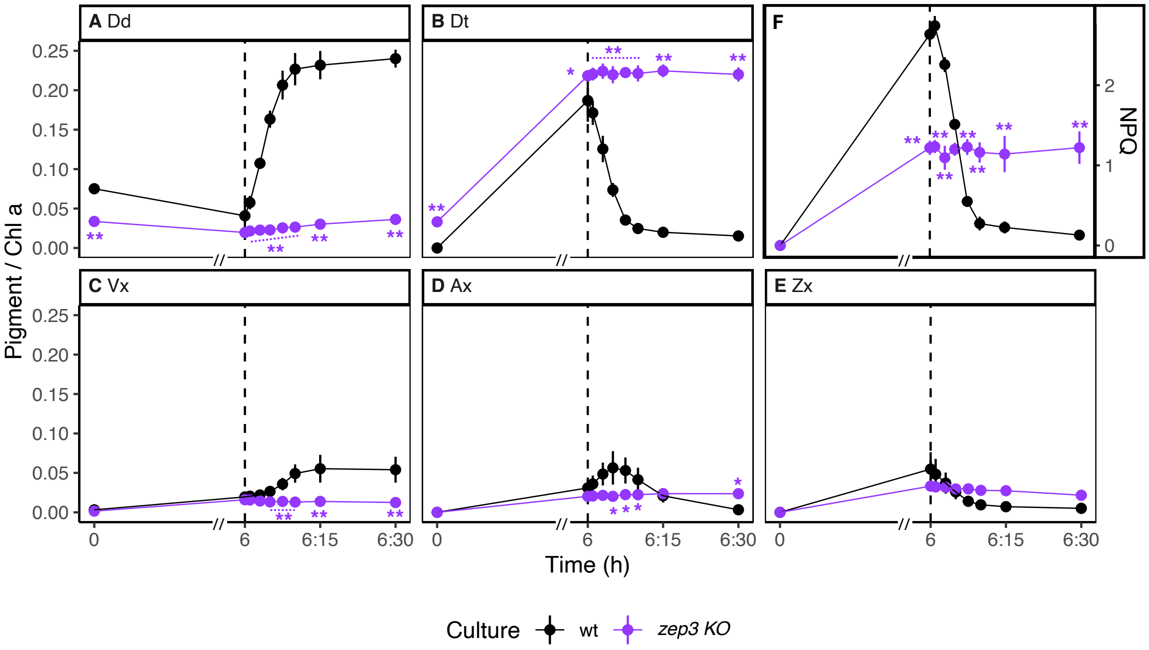


**Fig. S13: Pigment content of Dd and Vx cycle pigments coupled with NPQ analysis in wt and *zep3 KO* after 5 days of HL:LL regime (average ± sd, n=3).** Pigment content is expressed as pigment:chlorophyll *a* ratio (mol/mol). Time 0 is defined as the beginning of the HL phase (i.e. the sampling point in LL that immediately precedes the onset of HL on the fifth day of HL:LL treatment). The 6 h of HL phase and 30 min of recovery phase (LL) are separated with a dashed line. On the x-axis, recovery phase (from 6 to 6:30 h) was artificially enlarged to allow better visualization; axis break is indicated by a double dash (//). (*A*) diadinoxanthin; (*B*) diatoxanthin; (*C*) violaxanthin; (*D*) antheraxanthin; (*E*) zeaxanthiAs n; (*F*) NPQ; due to the substantial lack of Dt recovery in this mutant (as shown by the presence of Dt at time 0 and 6:30 h), the calculated NPQ depicted in this figure is most likely underestimated (measured Fm used to calculate NPQ is most likely lower compared to the real maximum fluorescence that would be measured in a fully relaxed state). Statistical significance marks indicate significant differences between the corresponding mutant line and wt at each time point, according to adjusted p-value of multiple comparison t-test (*: $p < 0.05$; **: $p < 0.005$).

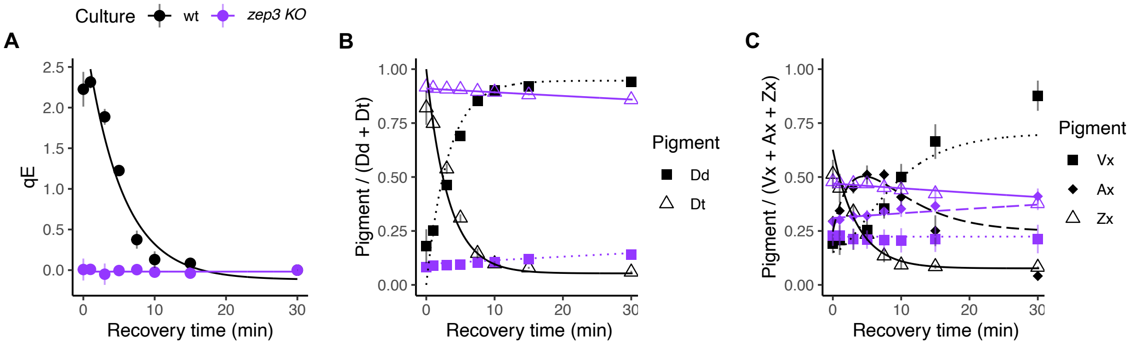


**D**

| | qE | Dt → Dd | Zx → Ax | Ax → Vx | Ratios | | | | | |
|---|---|---|---|---|---|---|---|---|---|---|
| | *R0* | $k_T$ [min$^{-1}$] | $k_Z$ [min$^{-1}$] | $k_A$ [min$^{-1}$] | $k_T/k_Z$ | $k_Z/k_T$ | $k_T/k_A$ | $k_A/k_T$ | $k_Z/k_A$ | $k_A/k_Z$ |
| **wt** | - 0.197 ± 0.025 | 0.310 ± 0.017 | 0.280 ± 0.035 | 0.167 ± 0.076 | 1.11 ± 0.08 | 0.90 ± 0.06 | 2.10 ± 0.81 | 0.53 ± 0.22 | 1.87 ± 0.61 | 0.58 ± 0.22 |
| ***zep3 KO*** | NA NA | 0.002 ± 0 | 0.005 ± 0 | 0.001 ± 0 | 0.4 ± 0 | 2.5 ± 0 | 3.33 ± 1.15 | 0.33 ± 0.14 | 8.33 ± 2.88 | 0.13 ± 0.06 |

**Fig. S14. First order kinetics of qE recovery and of re-epoxidation of xanthophyll cycle pigments in wt and *zep3 KO* during 30 min of recovery phase after HL (average ± sd, n=3).** (*A*) recovery rates of qE; (*B*) epoxidation of diadinoxanthin (Dd) to diatoxanthin (Dt); (*C*) epoxidation of zeaxanthin (Zx) via antheraxanthin (Ax) to violaxanthin (Vx); (*D*) corresponding rate constants of: qE recovery (R0); Dt to Dd epoxidation ($k_T$);, Zx to Ax epoxidation ($k_Z$); Ax to Vx epoxidation ($k_A$). For *zep3 KO*, no R0 values were determined as qE did not show significant changes during recovery. Due to the substantial lack of Dt recovery in this mutant (also shown in Fig. S11), the calculated qE is artificially always close to 0 (Fm'' measured at 30 min of recovery and used to calculate qE is almost equal to the Fm' of other time points sampled between 6 h of light stress and 30 min of recovery due to substantial lack of recovery). This doesn't mean that this mutant is not performing qE, but rather that this cannot be appropriately calculated due to substantial lack of recovery.

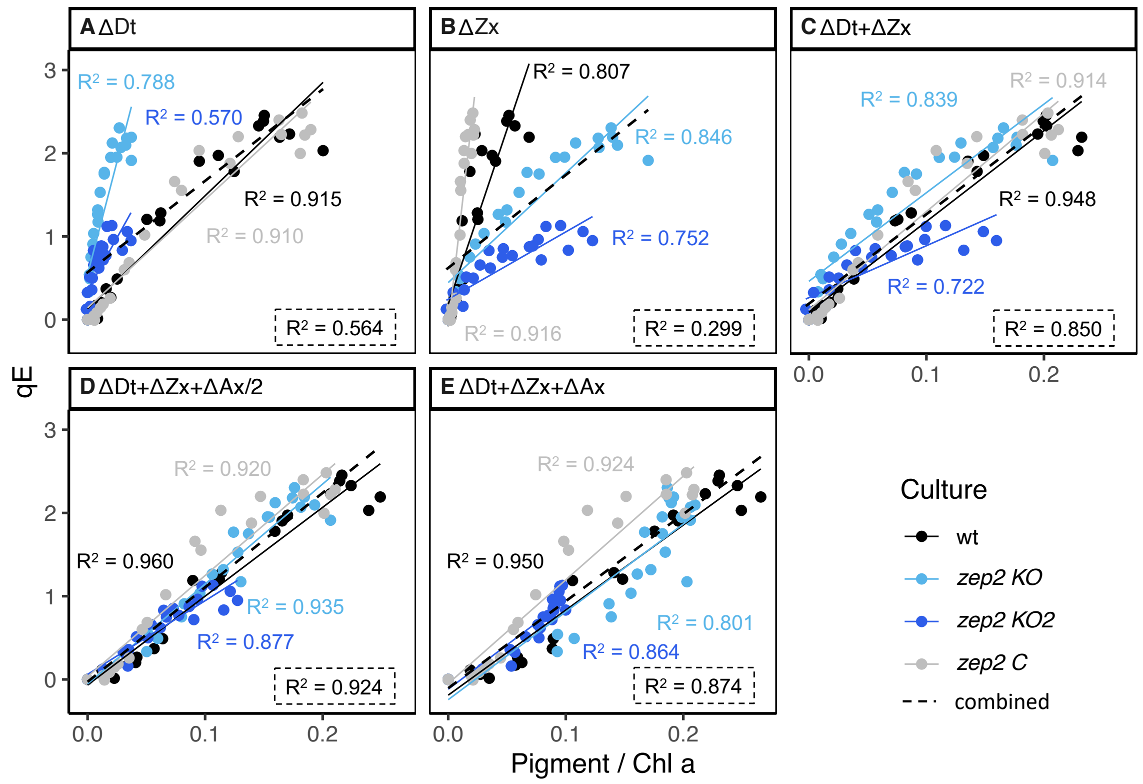


**Fig. S15. Correlation between qE and de-epoxidized pigments wt, *zep2 KO*, *zep2 KO2* and *zep2 C* during recovery from 6 h of HL, on the 5th day of HL:LL exposure.** Each point represents a separate measurement, repeated for 3 independent biological replicates. For each strain, different calculations of de-epoxidized pigment pool are displayed, with corresponding linear models and $R^2$. Dashed lines represent the linear model obtained when considering all points from different lines as one unique population (combined); corresponding $R^2$ is displayed in the dashed text box at the bottom right corner of each facet. (*A*) diadinoxanthin only; (*B*) zeaxanthin only; (*C*) diatoxanthin and zeaxanthin; (*D*) diatoxanthin, zeaxanthin and ½ antheraxanthin; (*E*) diatoxanthin, zeaxanthin and antheraxanthin.

**Table S1. Primer sequences used for mutant screening and PCR genotyping.**

| Mix name | Binding target | Primer F (5′-3′) | Primer R (5′-3′) | Size (bp) | T ann. (°C) |
|---|---|---|---|---|---|
| **PCR screening** | | | | | |
| *VDE* | *VDE* | VDE_longF: GTTCGATGTATTTCCCTGCC | VDE_longR: GCACCTTCTAGACCTTGCTC | 594 | 58 |
| *ZEP2* | *ZEP2* | ZEP2_longF: GGAATCCTTCTGTCACTAAC | ZEP2_longR: TGAAAGGTAAGTTTTCGTGT | 752 | 45 |
| *ZEP3* | *ZEP3* | ZEP3_longF: ATACGTAGCCACCACGGTGA | ZEP3_longR: GAAAACCGTGTACCCCGAGT | 1106 | 56 |
| **HRM** | | | | | |
| *VDE* | *VDE* | VDE_longF | VDE_shortR: CGGCTGATTCGGATCTTG | 151 | |
| *ZEP2* | *ZEP2* | ZEP2_shortF: CATAGAATCAATACAAGGCACAT | ZEP2_shortR: CGTGCTCGCAACGCGCGT | 111 | |
| *ZEP3* | *ZEP3* | ZEP3_shortF: CCGGCTCAAAGTGCTGATTG | ZEP3_shortR: GCGTTACTAGCGAGCTGGAT | 150 | |
| **PCR genotyping** | | | | | |
| *VDE* | wt *VDE* | VDE_1: GGCAAACTGAATTGGCG | VDE_A: TGGCTGGAAAAATTGCGTCC | 1791 | 58 |
| *vde KO* allele 1 | *vde KO* (allele 1) | VDE_2: GGCAAACTGAATTGGCC | VDE_A | 1791 | 58 |
| *vde KO* allele 2 | *vde KO* (allele 2) | VDE_3: TCTTCGGCAAACTGAATTAA | VDE_A | 1787 | 58 |
| *vde C* | *vde C* | VDE_1 | pPT_F: GTGACACTATAGAACCAGATCCC | 1409 | 58 |
| *ZEP2* | wt ZEP2 (variant for *zep2 KO* genotyping) | ZEP2_A: GACAAAACCTTGGCGAGCTG | ZEP2_1: CGCGCGTGGAGATACGG | 987 | 60 |
| *zep2 KO* allele 1 | *zep2 KO* (allele 1) | ZEP2_A | ZEP2_2: CGGAGGAACCGCCGTC | 968 | 60 |
| *zep2 KO* allele 2 | *zep2 KO* (allele 2) | ZEP2_B: GACCTACCCATTTTACCCGGT | ZEP2_3: TCAACATGCAGAGAGCGGTC | 140 | 60 |
| *zep2 C* | *zep2 C* | pPT_F | ZEP2_1 | 812 | 58 |
| *ZEP3* | wt ZEP3 | ZEP3_A: TTGGGTACGACGATTTCG | ZEP3_1: CCAGTACGGTAACATCCATG | 1948 | 60 |
| *zep3 KO* allele 1 | *zep3 KO* (allele 1) | ZEP3_longF | ZEP3_2: CCAGTACGGTAACATCCATC | 624 | 60 |
| *zep3 KO* allele 2 | *zep3 KO* (allele 2) | ZEP3_A | ZEP3_3: CCCGAGTAGGTGGCGG | 1693 | 60 |
| *zep3 C* | *zep3 C* | pPT_F | ZEP3_4: | 1374 | 60 |

| | | | | | |
|---|---|---|---|---|---|
| | | | CGGACGTTTGTTCCAAAACC | | |
| *ZEP2* (2) | wt ZEP2 (variant for *zep2 KO2* genotyping) | ZEP2_A | ZEP2_1b: GCGCGTGGAGATACGGA | 986 | 60 |
| *zep2 KO2* allele 1 | *zep2 KO2* (allele 1) | ZEP2_A | ZEP2_4: CAACGCGCGTGGAGAG | 980 | 58 |
| *zep2 KO2* allele 2 | *zep2 KO2* (allele 2) | ZEP2_A | ZEP2_5: CAACGCGCGTGGAGATACGA | 989 | 58 |

**Table S2. Average pigment content of all lines used in this study under pre-experimental conditions (low light control; average ± sd, n=3).**

| | Chl c | Fx | Fx-i | Vx | Dd | Ax | Dt | Zx | Chl a | β-car | c-β-car | TOT |
|---|---|---|---|---|---|---|---|---|---|---|---|---|
| rt (min) | 5.5 | 8 | 9.5 | 10.5 | 12 | 13 | 14.5 | 15.5 | 21.5 | 27 | 27.5 | |
| $\lambda_{max}$ (nm) | 337 445 632 | 448 | 441 | 418 441 470 | 447 477 | 447 475 | 453 481 | 453 479 | 414 430 663 | 454 480 | NA | |
| **wt** | 80.7 ± 24.9 | 269.4 ± 78.2 | 4.9 ± 3.7 | 1.5 ± 1.0 | 42.1 ± 14.6 | 0 | 0 | 0 | 554.2 ± 207.5 | 13.1 ± 3.8 | 0.7 ± 0.2 | 966.7 ± 332.0 |
| **ZEP2 KO** | 69.5 ± 20.9 | 237.6 ± 75.1 | 3.3 ± 2.4 | 1.2 ± 0.8 | 34.5 ± 12.6 | 2.6 ± 1.2 | 0 | 0 | 492.4 ± 163.6 | 11.3 ± 4.0 | 0.5 ± 0.5 | 852.9 ± 276.2 |
| **ZEP2 KO2** | 68.2 ± 7.8 | 225.8 ± 27.2 | 4.4 ± 1.1 | 2.6 ± 0.2 | 27.0 ± 3.4 | 5.5 ± 1.3 | 0 | 0.1 ± 0.2 | 443.7 ± 56.7 | 10.9 ± 0.9 | 0.9 ± 0.1 | 789.0 ± 97.9 |
| **ZEP3 KO** | 85.2 ± 9.7 | 274.9 ± 41.1 | 7.1 ± 3.7 | 1.6 ± 0.2 | 40.2 ± 6.1 | 0 | 1.3 ± 0.4 | 0 | 588.1 ± 76.3 | 13.7 ± 1.3 | 0.5 ± 0.04 | 1012 ± 131 |
| **VDE KO** | 84.8 ± 3.1 | 256.9 ± 15.6 | 10.3 ± 9.3 | 2.1 ± 1.3 | 45.7 ± 4.3 | 0 | 0 | 0 | 536.5 ± 32.0 | 16.4 ± 1.9 | 0.2 ± 0.3 | 979.8 ± 46.5 |
| **ZEP2 C** | 75.4 ± 6.9 | 268.0 ± 25.0 | 2.8 ± 2.0 | 1.6 ± 0.6 | 40.9 ± 4.1 | 0 | 0 | 0 | 538.7 ± 48.1 | 16.1 ± 1.8 | 0.8 ± 0.3 | 944.3 ± 88.2 |
| **ZEP3 C** | 94.2 ± 5.0 | 320.4 ± 1.9 | 4.6 ± 1.1 | 2.9 ± 0.3 | 35.0 ± 1.9 | 0 | 9.7 ± 0.9 | 0 | 635.5 ± 34.1 | 17.8 ± 1.5 | 0.3 ± 0.5 | 1120 ± 61.0 |
| **VDE C** | 70.4 ± 4.7 | 238.8 ± 10.2 | 2.2 ± 1.4 | 2.8 ± 0.3 | 40.1 ± 1.8 | 0 | 0 | 0 | 498.0 ± 30.0 | 15.0 ± 1.1 | 0.5 ± 0.5 | 867.8 ± 48.8 |

Pigment content is expressed as pmol:$10^6$ cells ratio (average ± sd, n=3; 0 = not detected). Representative retention time (rt) and $\lambda_{max}$ in our HPLC system are reported. Pigments are displayed according to retention time: Chl c: chlorophyll *c*; Fx: fucoxanthin; Fx-i: fucoxanthin isomer; Vx: violaxanthin; Dd: diadinoxanthin; Ax: antheraxanthin; Dt: diatoxanthin; Zx: zeaxanthin; Chl a: chlorophyll *a*; β-car: β,β-carotene; c-β-car: cis-β,β-carotene. Gray background highlights significant differences between the corresponding mutant line and wild type, according to adjusted p value of multiple comparison t-test ($p < 0.05$).

**Table S3. sgRNA coding-sequences used for CRISPR/Cas9 genome editing.**

| sgRNA | Target gene |
|---|---|
| GCGCGTGGAGATACGGAGAG | *ZEP2* |
| AGTACGGTAACATCCATGGT | *ZEP3* |
| CGGCAAACTGAATTGGCGGG | *VDE* |

**Table S4. Primer sequences used for generation and sequencing of complementation plasmids.**

| Sequence | Target(s) |
|---|---|
| TTTCTAATCACGATCGACCTGG | Amplification of *VDE* whole length (F) |
| GCACCGACCCAAAATCAAGAG | Amplification of *VDE* whole length (R) |
| AATGCGCTTTCGAACGGTG | Amplification of *ZEP2* whole length (F) |
| ACGTCGGCAACGCGAAG | Amplification of *ZEP2* whole length (R) |
| GGTAACCTTTCCTGGCCTTTCC | Amplification of *ZEP3* whole length (F) |
| AGCCTCTACACGTATGTGTTTCG | Amplification of *ZEP3* whole length (R) |
| GTGACACTATAGAACCAGATCCC | Sequencing of gene insertion site (F), **pPT_F** |
| TTAAGGAAGGATAGAGACT | Sequencing of gene insertion site (R) |
| CTCTCGGGCTCCAATTTTG | *VDE* intermediate sequencing (1) |
| GTTCGATGTATTTCCCTGCC | *VDE* intermediate sequencing (2), **VDE_longF** |
| CGGCTGATTCGGATCTTG | *VDE* intermediate sequencing (3), **VDE_shortR** |
| GCACCTTCTAGACCTTGCTC | *VDE* intermediate sequencing (4), **VDE_longR** |
| ATTTTACCCGGCAAGTAGC | *ZEP2* intermediate sequencing |
| ATGAAAAGATCTTGCAGTAT | *ZEP3* intermediate sequencing (1) |
| AGACAACCCACTACAACGGC | *ZEP3* intermediate sequencing (2) |
| TCTATGACACCCGTATAC | *ZEP3* intermediate sequencing (3) |
| GCTGAACCGAGCTCTTGTCC | *ZEP3* intermediate sequencing (4) |

Given the length of all target genes (*VDE*, *ZEP2*, *ZEP3*) several primers were used for sequencing of the complete gene inserted in the pPTbsr vector. Names of primers that were used also for mutant screening and genotyping (**Table S1.**) are indicated in bold.
